# In-silico evidence of non-operonic fusion transcripts in *Mycobacterium tuberculosis*: algorithm optimization and signatures of genome plasticity

**DOI:** 10.1101/2025.09.22.677293

**Authors:** Nikhil Bhalla

**Author notes:** **Correspondence should be addressed to:** Dr. Nikhil Bhalla (PhD, University of Delhi), Former Project Scientist – I/SRF, ICGEB, New Delhi, India., Former DBT-JRF/SRF at University of Delhi, New Delhi, India.

## Abstract

**Background:** The genome of *Mycobacterium tuberculosis* (Mtb) is known for its stable nature. It also contains transposases, redundant genes, repetitive DNA sequences, integrases, and remnants of lysogenized mycobacteriophages. These factors can result in intragenomic recombination, resulting in the formation of fusion transcripts. The present study aimed to identify signatures of long-distance gene fusion transcripts in RNA-seq data of clinical Mtb isolates.

**Methodology:** Three approaches based on separate principles (split read alignment, repurposing STAR chimera, and transcript de novo assembly). The intersections of fusion calls between the three approaches that showed maximum performance were used for detecting fusions with real RNA-seq datasets of Mtb.

**Results:** The junction of the split reads approach and the repurposed STAR chimera showed a high performance (F1 > 0.9). Sequence characteristics, clustering, and gene burden of operonic and long-distance gene fusions were consistent between the two independent real datasets, showing robustness of the optimized strategy. Fusion transcripts showed lineage specificity and signatures of indirect involvement of transposases, and transposition accessory genes (Rv1199c, Rv2512c, Rv3115, Rv0395, Rv2808, and Rv3327) in intragenomic recombination, resulting in the formation of fusion transcripts. The fusions mainly were within transposases, PPE, PE_PGRS family proteins, and some isolated fusions were of genes involved in the MoCo pathway, vesicle transport, and lipid turnover.

**Conclusions:** The observed fusions are likely driven by natural recombination, resulting in the formation of fusion proteins, coregulating proteins, or disruption. The study shows that the Mtb genome, especially of clinical isolates, may not be as stable as believed.

**Importance:** The Mtb genome is believed to be stable, clonal, and immune to HGT, and thus, only SNPs and INDELs are thought to drive evolution. However, the drastic differences in phenotypes such as growth kinetics, virulence, and metabolic rate observed in clinical isolates compared to laboratory strains cannot be entirely attributable to SNPs and INDELs. The Mtb genome contains transposases and other accessory genes that can drive intragenomic recombination, bringing distant genes closer. As a result, there is a possibility of the occurrence of fusion transcripts. Growing evidence and our previous contributions also suggest changes in gene repertoires and gene copy numbers, which are also likely driven by intragenomic recombination events. This study presents optimization of a robust and easy-to-implement fusion calling algorithm using traditional bioinformatic calls. Using the same, we report fusion transcripts of non-operonic genes in the RNA-seq data of clinical Mtb isolates.

## Introduction

Tuberculosis, a zoonotic and highly contagious disease older than humankind, is continuing to plague low to middle-income tropical nations, devastating economies at the level of nations, and at personal levels of infected patients and their families. With an estimated 10.8 million incidence in 2023 alone, Southeast Asian and African countries account for 69% of total burden (1). These continents harbour more than 50% of the world’s population, and the countries are poverty-ridden, often with poor healthcare infrastructure and availability. The causative agent, *Mycobacterium tuberculosis* (Mtb), has evolved into forms that are drug-resistant, drug-tolerant, and sometimes show erratic virulence in some patients leading to prolonged treatment and often fatalities. The laboratory strain of Mtb-H37Rv was stored in 1905 by Dr. Edward R Baldwin (2). This strain became a laboratory model to test drugs and understand Mtb biology in greater detail. Working with Mtb H37Rv is convenient as its doubling time and behavior in response to medication and other reagents are very predictable (3). These factors make its use convenient, but the when same experiments are carried out with clinical isolates, results often correlate poorly (4–7).

In the present-day clinical Mtb isolates, drastically long and differing doubling times, maximum CFU load in cultures at the stationary phase are well known (8). Differences in mechanisms of induction of immune response (clinical isolates were found to involve B-cell infiltration, rather than clinical isolates that induced T-cell infiltration) are reported (9). Upon phylogenetic clustering with various techniques, Mtb strains can be classified into lineages with some degree of association with certain factors. For example, lineage 1 is known to cause higher proinflammation, apoptosis, and moderate intracellular CFU burden compared to lineage 2, and other immunity-specific perturbations are discussed in detail by Chakraborty et al (8). These differences were well known; however, the Mtb genome is believed to be highly stable, clonal, and does not acquire new genes through horizontal gene transfer due to a thick cell wall that could mediate these differences (10,11).

Factors like variations in genomes between laboratory strains from different labs (12), gene repertoire differences (13,14), and the phenomena of presence-absence of the region of differences (15) are well known. The growth kinetics and clinical manifestation differences in lineages of Mtb are thought to be caused by SNPs in crucial loci, large sequence polymorphisms, presence/absence of regions of differences, and gene repertoire differences (13,14,16–19). In addition, Mycobacteriophages exist naturally, often in environments where Mtb thrives (20). These mycobacteriophages are not highly specific to Mtb, so they can carry foreign genes from other bacteria/species, and after attacking Mtb, may lysogenize, resulting in phage-driven lateral transfer (21,22). Therefore, despite HGT’s absence, Mtb is not immune to lateral transfer. Also, the Mtb genome contains many insertion sequences, transposons, and repetitive elements that can participate in intragenomic recombination events. These factors may cause fusion transcripts to occur in transcriptomic data. Given these factors, it becomes imperative to revisit a widely accepted understanding of the Mtb genome - that it is stable and clonal. The mentioned factors formed the basis of our study: the fusion of genomic loci can bring genes under the same promoter, thus resulting in the formation of fusion transcripts. In some cases, it can also result in multidomain fusion proteins, if translated. The objective of the present study was to determine whether fusion transcripts in high-throughput transcriptome data exist.

Currently, the published tools for identifying fusion transcripts are focused on eukaryotes and depend on the principle of alternative splicing and the presence of exons, both of which are absent in bacterial systems, including Mtb. The approaches to identify fusion transcripts in bacteria included extracting split reads from alignment data, *de novo* transcript assembly, mapping of contigs, and tweaking the eukaryotic-specific method. We report 1) a robust approach to determine fusion transcripts and the genes involved, 2) sequence characteristics of operonic and long-distance genes that fuse to form fusion transcripts, and 3) report high-confidence fusions found independently in two different datasets. The present study’s findings may help understand Mtb biology in greater detail, and they touch on the overlooked aspects in Mtb literature.

## Results

### Optimization strategy to determine high-confidence fusion transcript calls

Five differently seeded fusions (n=200 fusions/sample) were introduced in the CDS reference file of Mtb H37Rv. The details of the incorporated fusions were known and thus treated as ground truth. Short 150 bp x 2 PE reads (n=2M) were synthesised in-silico using the transcripts with fusions incorporated. These simulated reads were processed differently in the three approaches. Approach-1 included mapping with minimap2 to the reference genome and identification of reads that mapped to two different regions on the genome. Such reads are flagged as SA (Split-Alignments) reads in BAM files. The split reads were clustered, and fusion calls were determined. In approach 2, the STAR inbuilt option to determine chimeric transcripts was utilized, and junctions were converted to a format consistent with the output of other methods. The third approach included de novo assembly using Trinity assembler and mapping of the resulting contigs on the reference genome using minimap2. The rest of the steps were similar to the steps followed in approach-1. Once fusion calls were made, the calls were intersected with genomic CDS coordinates, and genes such as PE, PPE, IS6110, rRNA, and tRNA were flagged. The standardized calls were finally compared with the ground truth, and metrics like true positive, false positive, and false negative counts were determined. More conclusive metrics were determined using these counts: precision, recall, and F1 scores. A schematic representation of the optimization strategy followed is shown in Figure 1A.

**Figure 1.**
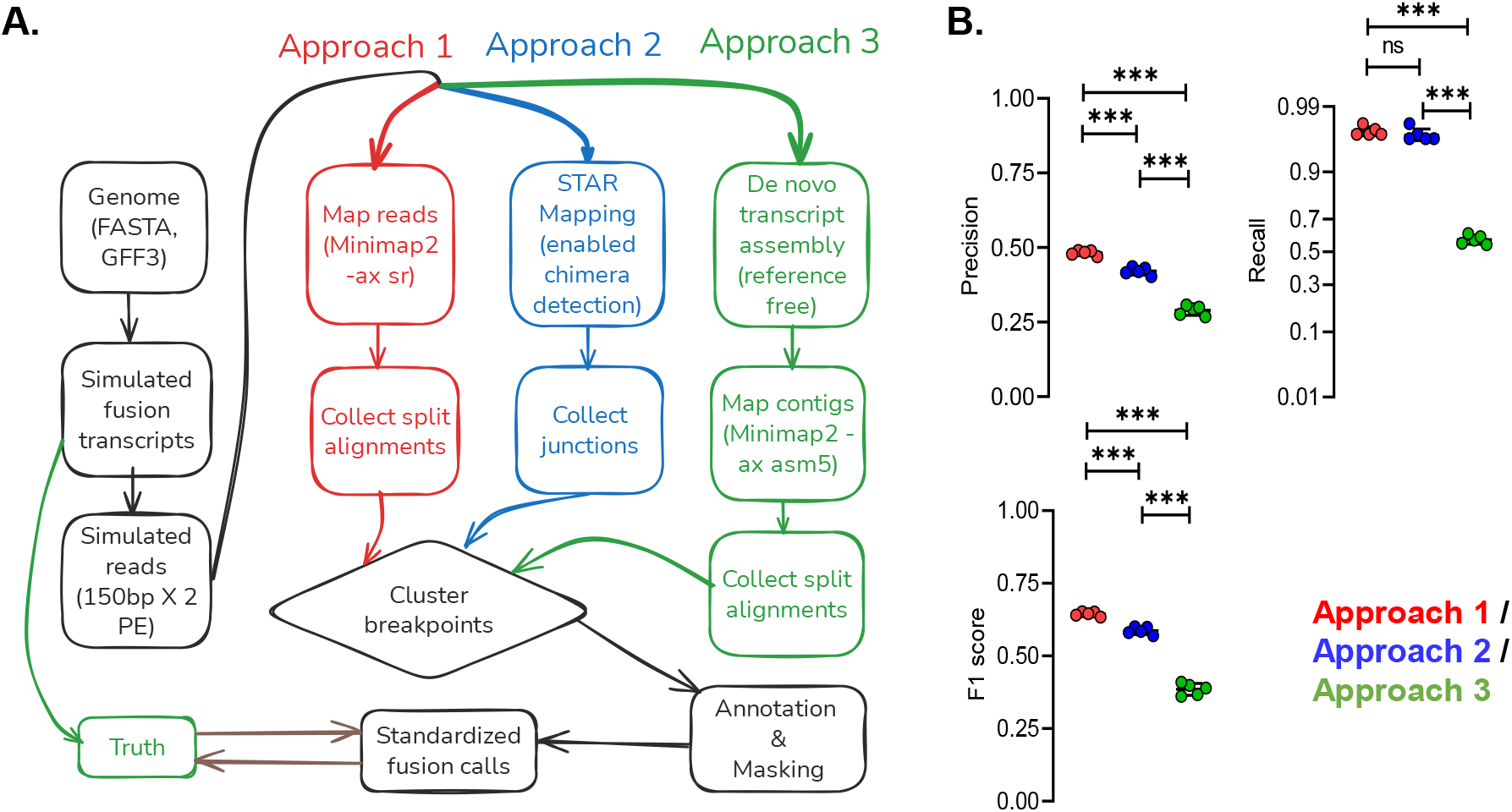
Independent approaches, their comparison, and optimization strategy to identify high-confidence fusion transcripts. The fusion transcripts were simulated using the CDS of Mtb H37Rv reference, and short reads were synthesized using WGSIM from the simulated transcript sequences. The simulated reads were processed using three approaches. Approach 1 included mapping with reference and using split alignment reads with respect to the reference genome. Approach 2 included using the STAR aligner with enabled chimeric transcript detection. The junctions produced were parsed subsequently. Approach 3 included de novo transcriptome assembly followed by the steps of Approach 1. The steps, such as clustering break points, annotation, and masking of known repetitive genes, were common in the three approaches. Finally, standardized fusion calls were compared with the ground reality of fusions in simulated transcriptome data. B: The precision (Equation 1), recall (Equation 2) and F1 scores (Equation 3) were determined for each approach independently.

### Independent approaches showed sub-optimal F1 scores

Approach-1 was based on identifying split reads after alignment to detect fusion transcripts, showed a precision value of 0.48 ± 0.01; a recall value of 0.97 ± 0.004; and an F1 score of 0.64 ± 0.008. approach – 2, which used the STAR-built tool to identify chimeric transcripts, showed a precision value of 0.42 ± 0.01; a recall value of 0.97 ± 0.01; and an F1 score of 0.58 ± 0.013. Approach–3, which was based on alignment of de novo assembled contigs to reference and then identifying split reads (in this case, contigs) after alignment to detect fusion transcripts, showed a precision value of 0.29 ± 0.02; a recall value of 0.58 ± 0.03; and an F1 score of 0.39 ± 0.02. Amongst the three approaches, the split read approach (approach-1) showed high values of performance metrics. However, the precision value was low, which decreased the overall F1 score (Figure 1B). The scores of individual replicates are presented in Supplementary Table 1.

### A union of selected intersections of fusion calls by the three approaches showed high metrics

The Venn sections were assigned letters A-G, as shown in Figure 2A. The fusion call numbers in each section were plotted and seemed within range. Sections A, B, C, D, E, F, and G contained 48.2 ± 7.3, 116.4 ± 9.8, 2.2 ± 1.6, 1.2 ± 1.1, 86.4 ± 6.5, 0.4 ± 0.5, and 112.8 ± 6.8, respectively. Counts were consistent with replicates made with different seeds (Figure 2A). Section-specific counts for each replicate subjected to different approaches are represented in Supplementary Figure 1. Similarly, metrics for each section were determined independently. Sections E and G showed high true positive counts: 82 ± 6.8 and 110.8 ± 6.72, respectively, and the rest showed near-zero accurate positive counts (Figure 2B). Sections A and B showed high false positive counts (a lower false positive count is better): 48.2 ± 7.3 and 116.4 ± 9.8, respectively, and the rest showed lower counts (C=1.6 ± 1.5; D=1.2 ± 1.1; E=4.4 ± 1.9; and G=2 ± 1.6) (Figure 2C). In case of false negatives (lower is better), sections A-D and G showed 199.8 ± 0.37 counts, while sections E and G showed 118 ± 6.8 and 89.2 ± 6.7 counts (Figure 2D). Since precision, recall, and F1 score are derived from the values of TP, FP, and FN, they give a better idea of performance. For these metrics, a high score is more desirable. In case of precision: sections A-B showed a cumulative score of 0 while section F scored 1 in one of the five replicates (the rest were 0). Sections E and G showed a cumulative precision score of 0.96 ± 0.02 (Figure 2E). In case of recall, sections A, B, and D showed zero scores in all replicates, while section C showed a score of 0.005 in 3 out of 5 seeded replicates, and section F1 showed a score of 0.005 in one out of 5 seeded replicates. Section E showed a F1 = 0.41 ± 0.34 score, and section G showed a F1 = 0.55 ± 0.34 (Figure 2F). The trend was similar in the case of F1 scores, with sections E (F1 = 0.57 ± 0.03) and G (F1 = 0.7 ± 0.02), outperforming other sections (Figure 2G). To select the sections that showed maximum performance, a checkerboard was used to determine better performing intersects. Sections E and G showed maximum scores and were found best. It is also worth mentioning that the number of false positives was low and thus acceptable in C, D, E, F and G sections (Figure 2C and 2H). At this point, sections E and G were found to perform with adequate, if not with the best, performance metrics. As a result, a union of sections E and G was scored against the ground truth, and the metric values surpassed our expectations. To recall, 200 fusions were randomly introduced in the simulated datasets. The true positives in the case of E∪G were 192.8 ± 1.92, false positives stood at 6.4 ± 3.5, and false negative counts were 7.2 ± 1.9 (Figure 2I). The precision, recall, and F1 scores in the E∪G fusion calls were found to be 0.96 ± 0.017, 0.96 ± 0.009, and 0.96 ± 0.01, respectively (Figure 2J). Based on these results, the fusion calls in E∪G were deemed high-confidence—the section-specific metric scores for each replicate separately in Supplementary Table 2. With the actual Mtb RNA-seq dataset, the fusion calls in E∪G have been determined and reported in subsequent sections.

**Figure 2.**
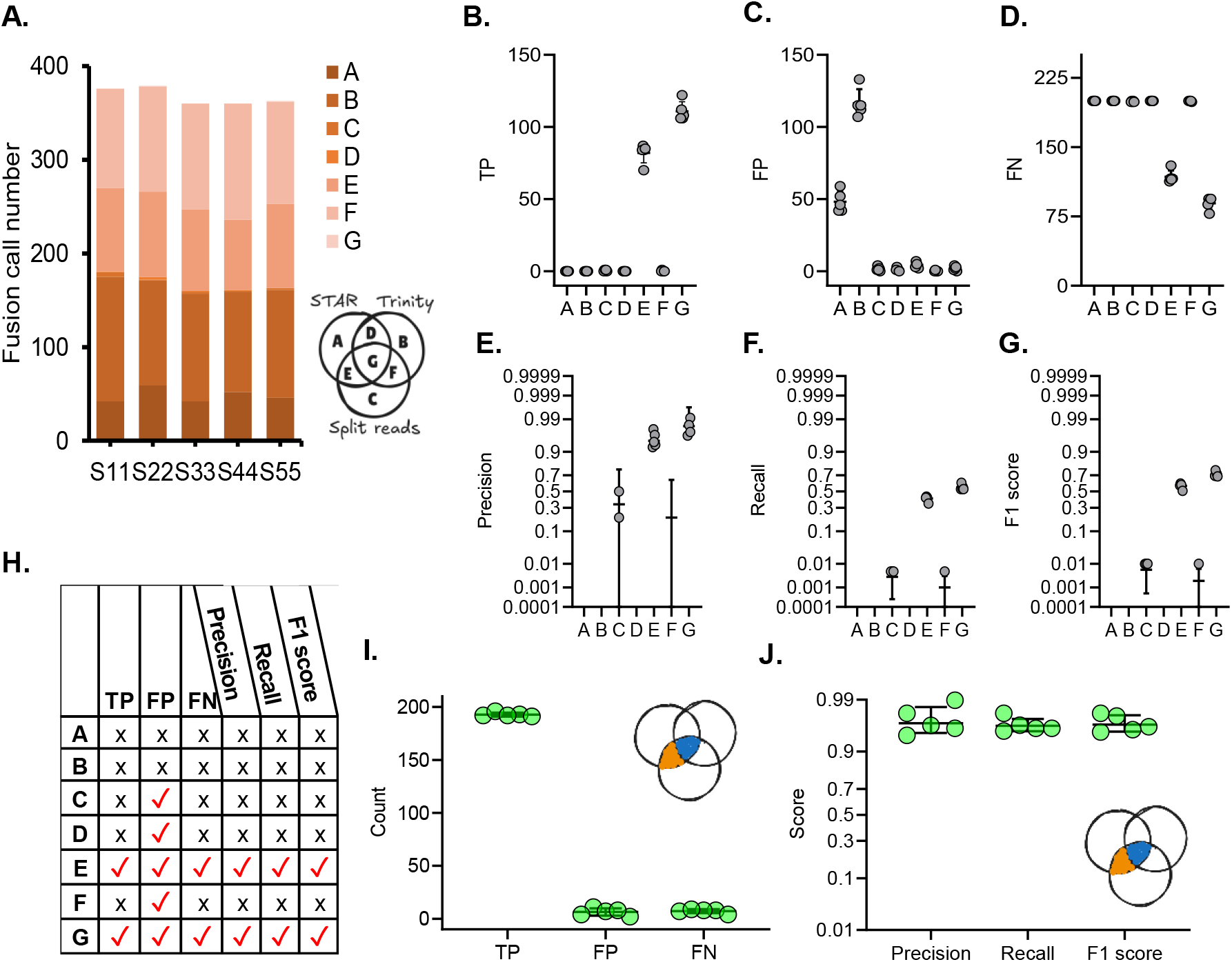
Selecting the intersections of fusion call sets with the best performance in terms of recall, precision, and F1 score. A: The fusion calls for each replicate (different seed) determined with the three approaches were subjected to Venn analysis independently (according to seeds). The number of calls in each intersection was determined, and an overview is presented. B: Fusion calls in each section were matched with ground truth, and metrics (TP, FP, FN, recall, precision, and F1 scores were determined. Each section of the intersection on the Venn diagram was assigned an alphabet from A to G. C: True positives (n = 200) in each section are shown. D: False positives in each section (lower is better). E: False negative in each section (lower is better). Figures E, F, and G show precision (high is better), recall (high is better), and F1 scores (high only if recall and precision both are high, high is better), respectively. H: Based on the metrics observed for each section, a summary was made in the form of a checkerboard. A tick mark (✓) indicates an acceptable metric score. From this, sections E and G were found to perform with maximum performance. I: As sections E and G were observed to have more acceptable metrics, the union of sections E and G was compared with the ground truth. This revealed a TP value close to the actual ground truth. J: The union of E and G sections, when compared with ground truth, showed >0.9 score of precision, recall, and F1 score.

### Fusion call metric patterns showed consistency across independent datasets

By this point, we had determined that the intersection of fusion calls made by approach-I (split reads based) and approach-II (STAR based) show a high F1 score based on the optimization done in the previous section using simulated datasets. The procured real datasets (RNA-seq data) were processed using approach-1 and -2, and the fusion calls in the intersection were determined for the two datasets. In dataset-1 (PRJEB29197) (23), n=153768 were identified exclusively by approach-1; n=778 were identified solely by approach-2, and both approaches identified n=799. With dataset-II (PRJEB8783) (24), n=180412 were detected solely by approach-1, n=6 exclusively by approach-II, and both approaches detected n=183. Approach-1 resulted in higher fusion counts while approach-2 consistently showed low fusion counts (Figure 3I/II-A). PLSDA analysis of transcript metrics resulted in diverging clusters of operonic and non-operonic transcripts. The operonic transcripts spread was higher than long-distance fusion calls, and the results were similar in both datasets (Figure 3I/II-B). Reads originating from genes with repetitive/redundant DNA can potentially misalign and give false positives. We determined k-mer (nt=8) dependent redundancy values and found higher sequence redundancy in fusions of operonic genes and lower redundancy in fusions of genes farther away, and in fusion transcripts. The results were statistically significant and similar in the two datasets (Figure 3I/II-C). Again, the binned count histogram and the Gaussian distribution showed higher sequence redundancy in operonic genes in fusion transcripts. The results were similar in both datasets (Figure 3 I/II-D). Gene link burden, the number of other genes with which it forms a fusion transcript, was determined in all pairs of intersecting sets and plotted. Most genes had fewer than 15 links in both datasets (Figure 4 I/II-E). The results indicated consistency in the sequence patterns of two independent patterns, implying the optimized approach gave robust and reproducible results with different datasets. The results also indicate fusions between genes separated by longer genomic distance were of lower redundancy, implying that misalignment may not have occurred that resulted in a long-distance gene fusion call.

**Figure 3.**
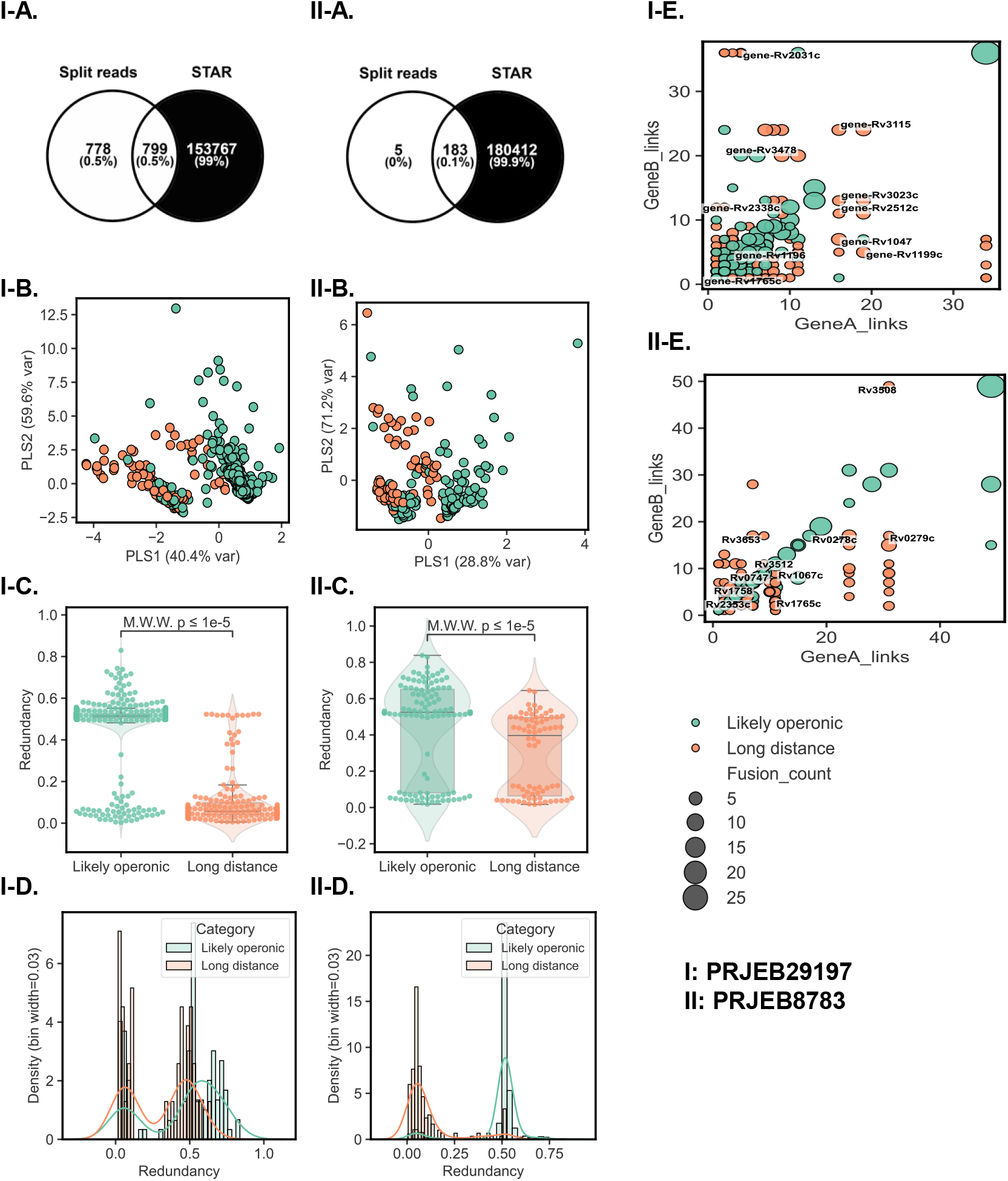
Fusion call metric patterns showed consistency across independent datasets. I/II-A: The intersection of fusion calls made by STAR and split-read approaches was determined. I/II-B: The fusion transcripts in the intersect were analyzed in terms of K-mer (of 8 nucleotides) based sequence redundancy, counts of gene A/B links, and the number of fusion isoform transcripts of the same pair. The values were scaled, and PLSDA analysis was carried out. I/II-C: The gene pairs were categorized as operonic if they were 6 or fewer genes apart, and redundancy scores were plotted. I/II-D: Density vs redundancy distribution. The redundancy range (0-1) was binned into 30 equal ratios. Fusion calls (of the intersection) were counted, and the density (Equation 4) within each bin was represented in the form of histograms. The curves show a smoothed Gaussian estimate trend of density vs redundancy. I/II-E: The gene count (for all genes in the intersection) in all fusion pairs was computed. The counts for each gene were plotted to represent the gene burden pattern. Abbreviations: M.W.W.: Mann– Whitney–Wilcoxon test. Note: Some dataset-1 samples were corrupted. They were repaired using repair.sh from bbtools before adapter trimming and subsequent processing.

**Figure 4.**
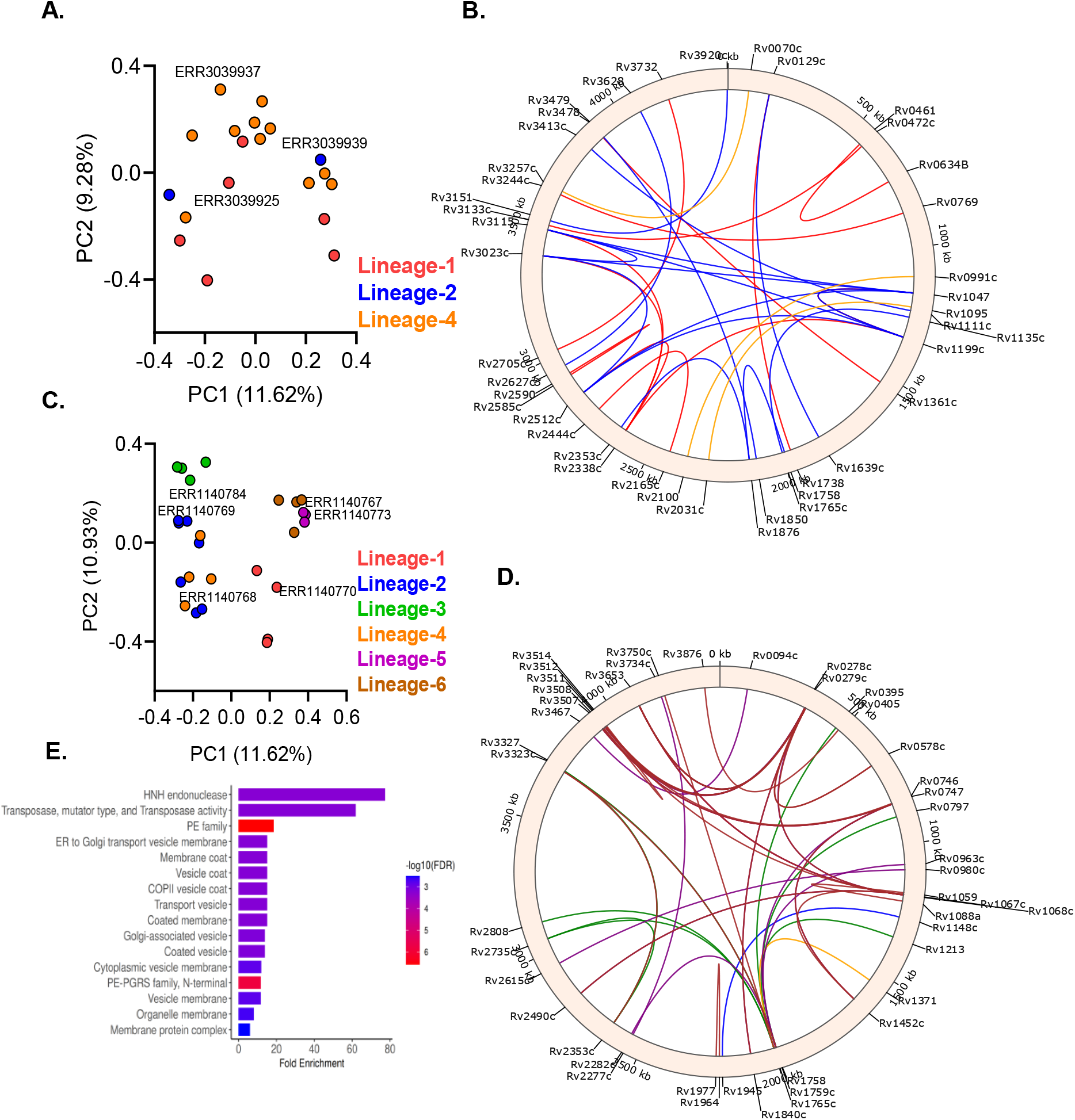
Long distance fusions were lineage showed signatures of lineage specificity. The intersection of fusion calls made by approach-1 and -2 for each sample was compiled into a binary of presence and absence, and principal coordinate analysis was performed. A: PCoA of dataset-1, C: PCoA of dataset-2. The fusion calls with 6 or fewer genes apart were removed, and a representative was marked on PCoA plots of each lineage, which were plotted in circos plots. B, D: Long distance fusion links identified in dataset-1 and -2, respectively. E: The genes taking part in long-distance fusions were compiled and subjected to gene ontology clustering, which revealed enrichment of various processes, with many that can potentially take part in recombination events. Note: The link colors in circos plots correspond to colors assigned to lineages in PCoA plots. The circos plots were made using the pycirclize Python library. Abbreviations: Principal coordinates.

### Long-distance fusions are numerous, mostly unique, and some common to separate datasets

The Total number of links observed in dataset-1 was 799 (non-operonic = 178; operonic n = 621), and in dataset-II was 183 (non-operonic = 76; operonic n = 107). The likely operonic links (transcript fusions) were removed, and the fusion calls were binned that lay in Rv3001-Rv0001-Rv1000 genes (bin-1), Rv1001-Rv2000 genes (bin-2), Rv2001-Rv3000 genes (bin-3). Fusions were visualized in the form of a circos plot. This binning was necessary for a cleaner representation of low (Supplementary Figure 2A), moderate (Supplementary Figure 2B), and long-distance (Supplementary Figure 2C) genes having a part in fusion transcripts. The genes that showed links in both datasets analysed were used for analysis with stringDB. The study revealed gene co-occurrence and co-expression relationships of Rv1765c with Rv1148c and Rv1945. These two genes were also correlated by gene co-occurrence, text mining, co-expression, and protein homology. These results aligned with our hypothesis and the resulting data that fusion transcripts can cause co-regulation of two distant genes. It is also worth noting that none of the genes with observed relations as per StringDB analysis are tandem on the Mtb genome (Supplementary Figure 2D). Fusion transcripts of PE and PPE proteins, which are known to have high sequence redundancy and multiple copies, may be fusion artefacts only (Supplementary Figure 2D right). The genes found in both datasets (labelled on circos diagrams) were subjected to gene ontology clustering, which showed enrichment of processes/domain/pathways such as HNH endonuclease, HNH nuclease, DUF22 protein domain, PPE, Endonuclease family, Response to host immune response, other organisms, and biotic stimulus. Of particular interest were enrichment of endonucleases, nucleases, and endonuclease family terms (Supplementary Figure 2E). These also aligned with our hypothesis that fusion transcripts may be driven by intragenomic recombination and/or phage attacks.

### Long-distance fusions showed signatures of lineage specificity

Upon stratification of fusion calls from each sample, the high-confidence fusion calls resulted in partially separate clustering on principal coordinate analysis of dataset-1 (Figure 4A) and dataset-2 (Figure 4C). Fusion calls from a representative sample from each lineage were plotted for dataset-1 and dataset-2. This revealed lineage-specific links between long-distance genes. (Figure 4B and 4D). Upon gene ontology clustering of genes taking part in long-distance fusions, processes relevant to recombination and genome modification activities were enriched. Of these, the interesting ones included: Transposase, mutator type, and Transposase activity, which included Rv1199c, Rv2512c, and Rv3115; Mostly uncharacterized, including. DDE transposase retroviral integrase sub-family, and Transposase, IS111A/IS1328/IS1533, N-terminal which included Rv0395, Rv2808, and Rv3327; Transposition which included Rv1199c, Rv2512c, Rv3115, and Rv3327; Mostly uncharacterized, incl. DDE transposase retroviral integrase sub-family, and Transposase, IS111A/IS1328/IS1533, N-terminal which included Rv0395, Rv2808, and Rv3327; Endonuclease activity which included Rv1148c, Rv1765c, and Rv1945; and DNA recombination which included Rv1199c, Rv2512c, Rv3115, and Rv3327. Other processes/terms that showed enrichment were PE, PE-PGRS, intracellular vesicle transport, vesicoal coat, and associated processes, Mixed, incl: decaprenyl diphosphate synthase-like, and Cell wall (Figure 4E). A complete enrichment table is provided in Supplementary Table 3. The results indicate involvement of recombination-relevant proteins as well as proteins that are essential for Mtb biology, virulence, and transport and processes associated with energy metabolism.

### Lineage specific long distance fusion transcript landscape is dominated by transposases, PPE and PE_PGRS family genes

Rv1199c (putative IS1081 transposase) was found to form fusion transcripts with Rv1047 (Probable transposase) in lineage 4, Rv2512 (Transposase for IS1081) in lineage 2, and Rv2338 (molybdopterin biosynthesis protein MoeW) in lineage 1 (P<0.001, positive association) and in lineage 4 (P<0.01, negative association). The fusion transcript of Rv1361c (PPE19) with Rv3478 (PPE60) showed positive and negative association in lineage 1 (P<0.01) and lineage 4 (P<0.05), respectively. Rv2338c also formed fusion transcripts with Rv2512c (Transposase for IS1081) and Rv3115 (probable transposase), both in lineage 1 (P<0.05, positive association). These fusion transcripts were observed in dataset-1 (Figure 4B, Table 1). Rv0278c (PE_PGRS3) formed fusions with Rv1068 (PE_PGRS20) (P<0.01, positive association), Rv1840c (PE_PGRS34) (P<0.01, positive association), and Rv3653 (PE_PGRS61) (P<0.05, positive association) in Lineage 5. Rv0279c (PE_PGRS4) fusion transcripts with Rv1840c (PE_PGRS34), and Rv3511 (PE_PGRS55) in lineage 5 (P<0.05, positive association in all) were observed. Rv0279c (PE_PGRS4) also formed fusions with Rv3511 (PE_PGRS55), Rv3512 (PE_PGRS56), and Rv3653 (PE_PGRS61) in lineage 6 (P<0.05, positive association in all). Rv0747 (PE_PGRS10) formed fusions with Rv1452c (PE_PGRS28) in lineages 5 and 6 (P<0.05, positive association). It also formed fusions with Rv1759c (wag22) in lineage 5 (P<0.05, positive association). Rv3653 (PE_PGRS61) formed fusions with Rv1068c (PE_PGRS20) in lineage one and lineage 5 (P<0.05, positive association). In lineage 2, the Rv3653-Rv1068c fusion was negatively associated (P<0.05). Rv1758 (Probable cutinase Cut1) formed fusions with Rv2282c (LysR family transcription regulator) (lineage 5, P<0.01, positive association), Rv3323c (MoaD-MoaE fusion protein MoaX) (lineage 6, P<0.01, positive association), and with Rv3327 (transposase fusion protein) (lineage 3, P<0.02, positive association). Rv1765c (HNH endonuclease) formed fusions with Rv2735c (hypothetical protein) and Rv2808 (hypothetical protein), and in lineage 3 (P<0.01, positive association). It also formed a fusion with Rv3850 (Conserved protein) in lineage 6 (P<0.02, positive association). A complete list of putative fusions is provided in Table 1 and visualized in Figure 4B and 4D.

**Table 1.**
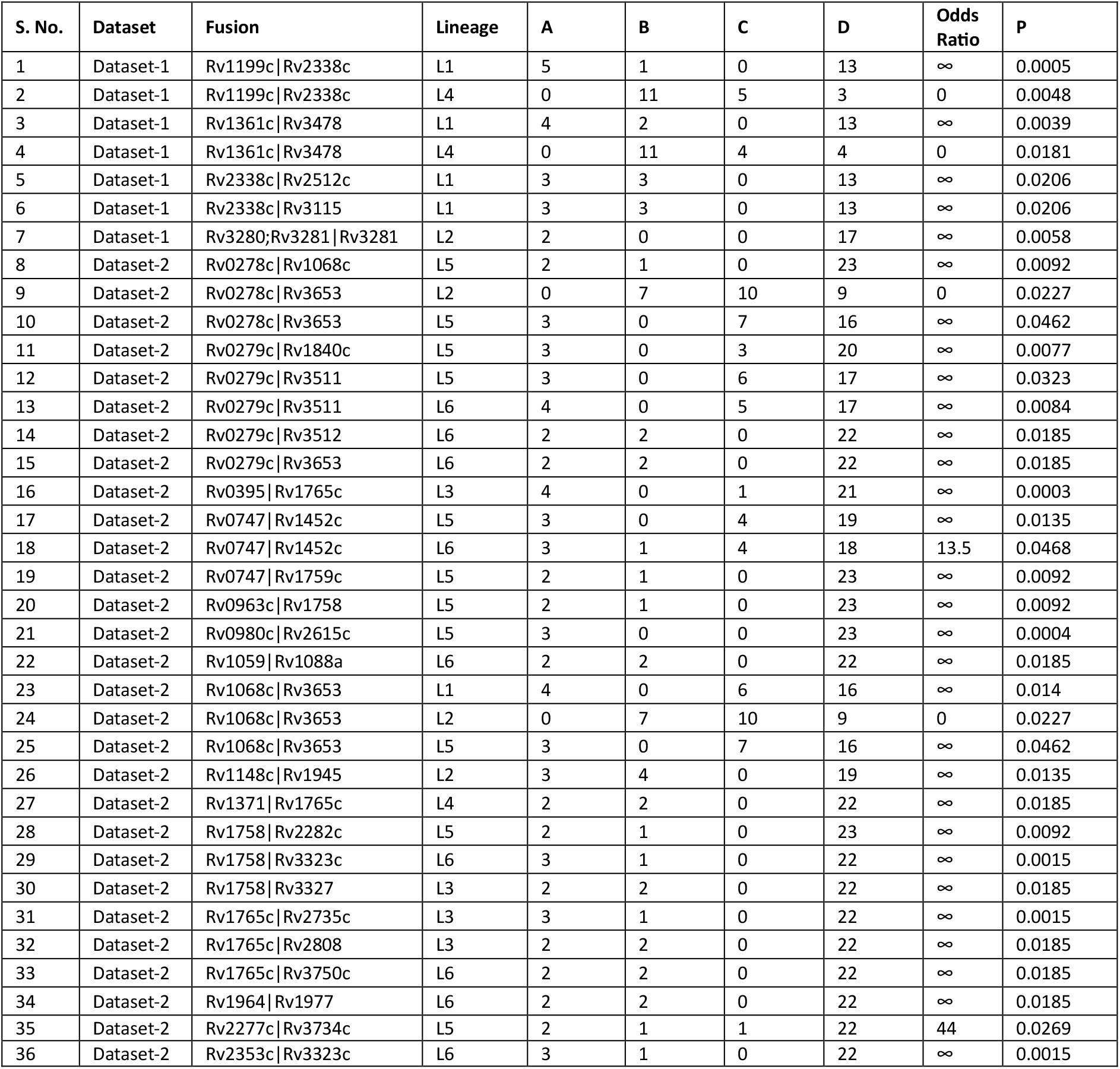
Association of fusions with lineages of Mtb. The fusion candidates that contained six or fewer local genes apart on the genome of Mtb H37Rv reference were removed. A two-tailed Fisher’s exact test was performed for association analysis between lineages and fusion transcripts. For each of the lineage’s number of samples of a specific lineage that have the fusion (count A), number of samples of a specific lineage that do not have the fusion (count B), number of samples not of the specific lineage that have the fusion (count C), number of samples not of the specific lineage that do not have the fusion were determined. The module fishers_exact from the scipy Python library was used. Odds ratio > 1 means positive association, = 1 means no association, and ≤ 1 means negative association of the fusion with specific lineages. Bioproject Dataset-1/2: PRJEB29197/PRJEB8783.

| S. No. | Dataset | Fusion | Lineage | A | B | C | D | Odds Ratio | P |
| --- | --- | --- | --- | --- | --- | --- | --- | --- | --- |
| 1 | Dataset-1 | Rv1199c\|Rv2338c | L1 | 5 | 1 | 0 | 13 | ∞ | 0.0005 |
| 2 | Dataset-1 | Rv1199c\|Rv2338c | L4 | 0 | 11 | 5 | 3 | 0 | 0.0048 |
| 3 | Dataset-1 | Rv1361c\|Rv3478 | L1 | 4 | 2 | 0 | 13 | ∞ | 0.0039 |
| 4 | Dataset-1 | Rv1361c\|Rv3478 | L4 | 0 | 11 | 4 | 4 | 0 | 0.0181 |
| 5 | Dataset-1 | Rv2338c\|Rv2512c | L1 | 3 | 3 | 0 | 13 | ∞ | 0.0206 |
| 6 | Dataset-1 | Rv2338c\|Rv3115 | L1 | 3 | 3 | 0 | 13 | ∞ | 0.0206 |
| 7 | Dataset-1 | Rv3280\|Rv3281\|Rv3281 | L2 | 2 | 0 | 0 | 17 | ∞ | 0.0058 |
| 8 | Dataset-2 | Rv0278c\|Rv1068c | L5 | 2 | 1 | 0 | 23 | ∞ | 0.0092 |
| 9 | Dataset-2 | Rv0278c\|Rv3653 | L2 | 0 | 7 | 10 | 9 | 0 | 0.0227 |
| 10 | Dataset-2 | Rv0278c\|Rv3653 | L5 | 3 | 0 | 7 | 16 | ∞ | 0.0462 |
| 11 | Dataset-2 | Rv0279c\|Rv1840c | L5 | 3 | 0 | 3 | 20 | ∞ | 0.0077 |
| 12 | Dataset-2 | Rv0279c\|Rv3511 | L5 | 3 | 0 | 6 | 17 | ∞ | 0.0323 |
| 13 | Dataset-2 | Rv0279c\|Rv3511 | L6 | 4 | 0 | 5 | 17 | ∞ | 0.0084 |
| 14 | Dataset-2 | Rv0279c\|Rv3512 | L6 | 2 | 2 | 0 | 22 | ∞ | 0.0185 |
| 15 | Dataset-2 | Rv0279c\|Rv3653 | L6 | 2 | 2 | 0 | 22 | ∞ | 0.0185 |
| 16 | Dataset-2 | Rv0395\|Rv1765c | L3 | 4 | 0 | 1 | 21 | ∞ | 0.0003 |
| 17 | Dataset-2 | Rv0747\|Rv1452c | L5 | 3 | 0 | 4 | 19 | ∞ | 0.0135 |
| 18 | Dataset-2 | Rv0747\|Rv1452c | L6 | 3 | 1 | 4 | 18 | 13.5 | 0.0468 |
| 19 | Dataset-2 | Rv0747\|Rv1759c | L5 | 2 | 1 | 0 | 23 | ∞ | 0.0092 |
| 20 | Dataset-2 | Rv0963c\|Rv1758 | L5 | 2 | 1 | 0 | 23 | ∞ | 0.0092 |
| 21 | Dataset-2 | Rv0980c\|Rv2615c | L5 | 3 | 0 | 0 | 23 | ∞ | 0.0004 |
| 22 | Dataset-2 | Rv1059\|Rv1088a | L6 | 2 | 2 | 0 | 22 | ∞ | 0.0185 |
| 23 | Dataset-2 | Rv1068c\|Rv3653 | L1 | 4 | 0 | 6 | 16 | ∞ | 0.014 |
| 24 | Dataset-2 | Rv1068c\|Rv3653 | L2 | 0 | 7 | 10 | 9 | 0 | 0.0227 |
| 25 | Dataset-2 | Rv1068c\|Rv3653 | L5 | 3 | 0 | 7 | 16 | ∞ | 0.0462 |
| 26 | Dataset-2 | Rv1148c\|Rv1945 | L2 | 3 | 4 | 0 | 19 | ∞ | 0.0135 |
| 27 | Dataset-2 | Rv1371\|Rv1765c | L4 | 2 | 2 | 0 | 22 | ∞ | 0.0185 |
| 28 | Dataset-2 | Rv1758\|Rv2282c | L5 | 2 | 1 | 0 | 23 | ∞ | 0.0092 |
| 29 | Dataset-2 | Rv1758\|Rv3323c | L6 | 3 | 1 | 0 | 22 | ∞ | 0.0015 |
| 30 | Dataset-2 | Rv1758\|Rv3327 | L3 | 2 | 2 | 0 | 22 | ∞ | 0.0185 |
| 31 | Dataset-2 | Rv1765c\|Rv2735c | L3 | 3 | 1 | 0 | 22 | ∞ | 0.0015 |
| 32 | Dataset-2 | Rv1765c\|Rv2808 | L3 | 2 | 2 | 0 | 22 | ∞ | 0.0185 |
| 33 | Dataset-2 | Rv1765c\|Rv3750c | L6 | 2 | 2 | 0 | 22 | ∞ | 0.0185 |
| 34 | Dataset-2 | Rv1964\|Rv1977 | L6 | 2 | 2 | 0 | 22 | ∞ | 0.0185 |
| 35 | Dataset-2 | Rv2277c\|Rv3734c | L5 | 2 | 1 | 1 | 22 | 44 | 0.0269 |
| 36 | Dataset-2 | Rv2353c\|Rv3323c | L6 | 3 | 1 | 0 | 22 | ∞ | 0.0015 |

## Discussion

The Mtb genome is not immune to intragenomic recombinations and mycobacteriophage attacks. These can cause the formation of fusion transcripts that may play specific roles in Mtb biology, especially of the clinical strains. The conventional tools to detect fusion transcripts are made for eukaryotes and rely on the presence of introns and alternate splicing, viz., StringTie (25), Arriba (26), FusionCatcher (27), and STAR-fusion (28). The NF-core/RNAfusion pipeline also relies on these tools and thus is not suitable for detecting fusions in bacteria (29). Thus, their direct implementation on bacterial data would not have been appropriate. Therefore, we had to benchmark multiple approaches and compare and select the intersection that showed maximum precision, recall, and F1 scores. Described in previous sections, the split reads approach for identification was inspired by the contributions of Ha et al (30) and Maher et al (31). The STAR-fusion approach was tweaked by making a GTF file containing fake exon coordinates. The command was also tweaked by optimizing parameters to be able to determine fusion transcripts (28). Trinity, a state-of-the-art tool, can build a full-length transcriptome without a reference sequence de novo. In theory, this ability makes it an ideal tool to assemble fusion transcripts, if any (32). Approach-1 was the fastest to execute, followed by Approach-2 and Approach-3, which used Trinity, and was too slow. In the end, the intersection of all three approaches were determined and the intersection of fusion calls with the two approaches-split reads (approach-I) and STAR (approach-II) were found to have high confidence. So, using trinity-based method (approach-3) was deemed unnecessary, which made the exercise time-saving and computationally resourceful. We observed that the optimized strategy for fusion transcript detection is robust and may be tested with other organisms and/or larger and more diverse simulated datasets.

With actual RNA-seq datasets, the observed consistencies of metrics between independent biological datasets, such as higher fusion calls by Approach-2 than Approach-1, higher overall spread of operonic fusions on PLSDA, statistically significant higher redundancy scores in likely operonic compared to long distance fusions, and more operonic fusions than long distance (which is expected also) demonstrate reproducibility and robustness of the optimized approach. Also, Approach-1 appeared more conservative than Approach-2 as it identified fewer fusion calls. This also indicates that each method captures a different kind of fusion. While some true fusions could exist in the unique non-overlapping sets also, the intersect was considered a high-confidence fusion call set. Both approaches converging to similar fusion calls and biological interpretations also add to the robustness of the strategy optimized. It is well known that redundancy in sequences can make precise alignment with short reads difficult (33–35). Therefore, the redundancy with genes can also cause false-positive fusion calls. Interestingly, lower k-mer redundancy in long-distance genes fusing in transcripts compared to likely operonic fusions was observed. The K-mer based redundancy metric was also based on the assumption that fusion transcripts are often of the genes with higher similarity such as rrs-rrl, groel(s), PPE, PE_PGRS etc. So, fusion transcripts of distant genes would have lower k-mer redundancy. Our observation of lower k-mer sequence redundancy in long distance fusion transcripts compared to likely operonic transcripts aligned with our assumption. These observations further strengthens that the long-distance fusions observed could indeed be valid. Also, the observation was consistent between the two datasets. This also suggests that the identified fusion calls, especially the long-distance ones, may not be sequencing artefacts. The Mtb genome is reported to have 749 operons, which is 54.5% of the entire Mtb genome, and our findings of higher gene link burdens of likely operonic gene transcript fusions corroborate (36). Like other metrics, this observation was also consistent in the two independent datasets.

One of the gene families known to be highly redundant in Mtb includes insertion sequences and genes that encode transposases for specific insertion sequences that upon transposition, disrupt the transcription of other genes (37,38). In our study, we have observed fusion transcripts of Rv1199c insertion sequences (IS1081) forming fusion transcripts with various other genes, such as moeW, and even with other insertion sequences, such as those encoded by Rv2512. MoeW tends to get impacted with more than insertion sequences encoded by multiple genes (Rv1199c, Rv2512c, and Rv3115). MoeW, a protein required for molybdenum cofactor biosynthesis, is essential for energy stress, virulence, and intracellular survival (39– 41). The observed fusion transcripts with insertion sequences suggest that the locus is more dynamic and may even be in the process of duplication and disruption. The PE_PGRS gene family encodes surface antigens with a conserved structural domain and depends on the ESX-5 system for secretion. These proteins are also known for redundancy in Mtb genomes (42,43). We have observed multiple fusion transcripts of PPE19-PPE60, PR_PGRS3-PE_PGRS34, etc. Multiple fusion transcripts of PE_PGRS and PPE genes indicate ongoing evolutionary processes. Since these are involved in virulence, the fusions may have a role in hyper-virulence observed in specific lineages. Gallant et al identified fusion transcripts of PPE38-71, which are two genes apart, and we have observed fusion of Rv2353c (PPE39), which is right next to the PPE38 on the Mtb reference genome (44). In our case, PPE39 formed a fusion with Rv3323 (moaX). The translocation of PPE39 to the site of moaX probably caused the formation of the previously reported fusion transcript of PPE38-71. Cutinase, encoded by Cut1 (Rv1758), catalyzes the cleavage of the ester bond of cutin lipids, and it was found to form multiple fusions with other proteins (45). Theoretically, Cut1-LysR fusion may form a lipid-sensing transcriptional regulator if fused in frame or cause co-regulation of both proteins; Cut1-MoaX fusion may couple the MoCo pathway with lipid catabolism through coregulation or disruption, and the presence of transposase in the transcript, with a part of Cut1, indicates an ongoing random transposition event. In addition to insertion sequences, PE_PGRS and PPE, and Cut1 forming fusions, we have also observed other proteins encoding endonucleases forming fusions with other proteins with unknown function, such as Rv1765-Rv2735c, Rv1765-Rv2808, and Rv1765-Rv3850.

Our hypothesis was based on insertion sequences, repetitive elements, and other genes that can cause intragenomic recombination in Mtb. It thus may bring distant genes closer in newer Mtb clinical strains. We observed the involvement of insertion sequences, repetitive elements, and transposases forming fusion transcripts with other transcripts encoding a completely different protein or proteins of their own families. The present study sheds light on a whole-genome-scale intragenomic recombination-driven formation of fusion transcripts in a lineage-specific manner. Whilst we have determined fusions, the exact structure or sequence of fusion transcripts warrants further investigation, perhaps by salvaging existing RNA-seq data or long-read WGS/RNA-seq. Also, the fusions observed may be attributable to multiple fusion transcript isoforms. This also warrants the need for additional sequencing experiments with clinical isolates. The present study provides enough computational evidence of fusion transcripts that are most likely driven by recombination events. Also, the findings may have a role in imparting lineage-specific phenotypes such as growth kinetic differences and erratic drug tolerance profiles. The present study also indicates that the Mtb genome may not be as stable as initially thought and provides a basis for further investigation on Mtb genome dynamics with a more holistic view, emphasizing recent advances in the field.

## Methodology

### Fusion transcript dataset preparation

The Mtb H37Rv (RefSeq: NC_000962.3) GFF3 and FASTA files were used for preparing fusion transcripts. GFFread (v0.12.7) was used to extract CDS from the RefSeq dataset. CDS of >300 bp were filtered and randomly selected based on the value of seeds specified (seeds = 11, 22, 33, 44, 55). The selected CDS to be fused were cut 120 nucleotides from the end (left CDS) and head (right CDS). The cut was done to retain codons in-frame in 60% of the fusions. The resulting fusions were concatenated with the original CDS extracted from GFF3 and genomic fasta. Python libraries: random, collections.namedtuple, Bio.SeqIO, and Bio.seq.seq biopython were used for preparing the simulated dataset (Supplementary 2.1). Using the concatenated CDS plus fusion transcript files as input, reads were simulated using WGSIM (v1.19.2) with specific options (e=0, d=250, s=30, N=200000, -1/-2 = 150) enabled.

### Determining fusion transcripts with Approach-I

The GFF3 file was converted to a bed file for CDS. CDS encoding rRNA, tRNA, IS6110, PE, and PPE were also compiled as a bed file for masking. The reads (simulated or adapter trimmed) were aligned to the Mtb H37Rv CDS dataset using minimap2 (v2.1). The alignment files were sorted using Samtools (v1.19.2). Pysam Python library parsed BAM, and bedtools as a linux subprocess to determine interval intersections. A window of 20 nucleotides was used to scan BAM for reads with SA tags (Supplementary 2.2). The junctions were subsequently annotated with the CDS bedfile created earlier. The resulting output was a bed file containing information on fusion transcripts, coordinates, and masking status (Supplementary 2.5).

### Determination of fusion transcripts with Approach-II

The CDS in the GFF3 file was reformatted into a GTF file, and the CDS were treated as exons. This was done using the awk command. The genomic fasta file and the produced fake GTF file were used as input for making the STAR (v2.7.11b) index. The reads (simulated or adapter trimmed) were aligned with STAR using the index created in previous steps with parameters that disabled intron-based splicing (--alignIntronMax 1) and enforced stringent chimeric detection settings (--chimOutType Junctions WithinBAM HardClip, --chimSegmentMin 10, --chimJunctionOverhangMin 10). The resulting Chimeric.out.junction files were reformatted using awk and collapsed into junctions. Supporting read counts of 3 were used, and then the breakpoint ends were slightly extended (2 left and three on the right nucleotides) to make slopped BED intervals. The sloped bed intervals were intersected with CDS coordinates using Bedtools (v2.31.1) intersect. This was done to annotate gene identifiers (Supplementary 2.3). The remaining fusion calls were compared with other methods.

### Determination of fusion transcripts with Approach III

The reads (simulated or adapter-trimmed) were de novo assembled into transcript contigs using Trinity (v2.9.1). The resulting contigs were aligned to the reference CDS using Minimap2 with the asm2 parameter. The alignment was sorted using the samtools sort module. Pysam Python library was used to parse the alignment BAM files to extract reads containing “SA” tags. The main alignment and alternative alignment positions were determined. The breakpoints in the alignment of SA reads were grouped into bins of 15, so slight variations in alignment positions of multiple reads could be clubbed together. Each read pair was treated as a possible fusion junction, and the average position was determined. The breakpoints were extended ±10 bp to catch overlapping regions, and the genomic reference bed file was used for intersecting and annotation (Supplementary 2.4).

### RNA-seq data acquisition and processing for the determination of actual fusion transcripts

A Literature search was carried out to filter out studies in which RNA-seq of cultured clinical isolates is done. Bioprojects PRJEB29197 and PRJEB9763 were identified, and the SRA datasets were downloaded using the European Nucleotide Database (ENA). The reads were adapter-trimmed using fastp (v0.23.4). The trimmed reads were subjected to the three approaches explained above. Metadata and corresponding articles contained information on lineages and drug resistance profiles. These were used to group samples for subsequent statistical analysis.

### Statistical analysis

Simulated reads were made with different seeds to optimize the fusion calling approach, and the gene transcripts were randomly fused. For each seeded dataset, a file of ground truth containing fusions used to make fusion transcripts was available. This ground truth file and the fusion calls determined with the three approaches were matched, and parameters like True positives (TP), True negatives (TN), and False positives (FP) were counted. Using these counts, metrics like precision, recall, and F1 score were determined (Supplementary 2.6). 2-tailed fisher’s exact test was performed for determining the association of fusions with specific lineages and those with P<0.05 and long distance genes were considered statistically significant and meaningful fusion transcripts.

### Formulae used

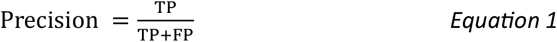

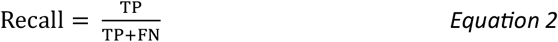

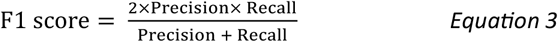

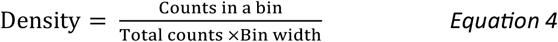

### Data visualization

Excalidraw was used to draw flowchart schematics and representative Venn figures. Venn analysis and diagrams were made in R using the ggVennDiagram library. Plots were made in Graphpad Prism. MS-Excel was used for data parsing and data representation. Pycirclize was used for making circos diagrams.

### Computational resources

All computational tasks were carried out on WSL2 running on an I5-8265U with 24 GB RAM on a consumer-grade personal computer, and more demanding tasks, such as adapter trimming and Trinity de novo assembly, were carried out on WSL2 on an AMD 5600G with 64GB RAM, a consumer-grade personal system. Wherever necessary, GNU Parallel was used to speed up processing.

## Supporting information

Supplementary 1 (Codes)

## Declarations

## Acknowledgement

The open-source tool developers and submitters of Mtb WGS in public repositories are acknowledged.

## Contributions

NB contributed to conceptualization, data curation, formal analysis, investigation, methodology, resources, visualization, and writing – original draft.

## Data availability statement

All data generated is contained within the manuscript and associated supplementary files.

## References

1. Global Tuberculosis Report 2024. 1st ed. Geneva: World Health Organization; 2024. 1 p.

2. Steenken W, Gardner LU. History of H37 strain of tubercle bacillus. Am Rev Tuberc. 1946 July;54:62–6.

3. Lakshmi R, Kumar V, Rahman F, Ramachandran R. Consistency of standard laboratory strain Mycobacterium tuberculosis H37Rv with ethionamide susceptibility testing. Indian J Med Res. 2012;135(5):672–4.

4. Devasundaram S, Raja A. Variable transcriptional adaptation between the laboratory (H37Rv) and clinical strains (S7 and S10) of Mycobacterium tuberculosis under hypoxia. Infect Genet Evol J Mol Epidemiol Evol Genet Infect Dis. 2016 June;40:21–8.

5. O’Toole RF, Gautam SS. Limitations of the Mycobacterium tuberculosis reference genome H37Rv in the detection of virulence-related loci. Genomics. 2017 Oct 1;109(5):471–4.

6. Romagnoli A, Petruccioli E, Palucci I, Camassa S, Carata E, Petrone L, et al. Clinical isolates of the modern Mycobacterium tuberculosis lineage 4 evade host defense in human macrophages through eluding IL-1β-induced autophagy. Cell Death Dis. 2018 May 24;9(6):624.

7. Jhingan GD, Kumari S, Jamwal SV, Kalam H, Arora D, Jain N, et al. Comparative Proteomic Analyses of Avirulent, Virulent, and Clinical Strains of Mycobacterium tuberculosis Identify Strain-specific Patterns*. J Biol Chem. 2016 July 1;291(27):14257–73.

8. Hailemariam TG, Tilahun M, Atnafu A, Gelanew T, Gebresilase TT, Tola MA, et al. Comparative growth kinetics and drug susceptibility of Mycobacterium tuberculosis lineages prevalent in Ethiopia: implications for tuberculosis treatment and management. Front Microbiol. 2025 Feb 4;15:1512580.

9. Mourik BC, de Steenwinkel JEM, de Knegt GJ, Huizinga R, Verbon A, Ottenhoff THM, et al. Mycobacterium tuberculosis clinical isolates of the Beijing and East-African Indian lineage induce fundamentally different host responses in mice compared to H37Rv. Sci Rep. 2019 Dec 27;9(1):19922.

10. Koleske BN, Jacobs WR, Bishai WR. The Mycobacterium tuberculosis genome at 25 years: lessons and lingering questions. J Clin Invest. 133(19):e173156.

11. Dos Vultos T, Mestre O, Rauzier J, Golec M, Rastogi N, Rasolofo V, et al. Evolution and Diversity of Clonal Bacteria: The Paradigm of Mycobacterium tuberculosis. PLoS ONE. 2008 Feb 6;3(2):e1538.

12. Ioerger TR, Feng Y, Ganesula K, Chen X, Dobos KM, Fortune S, et al. Variation among Genome Sequences of H37Rv Strains of Mycobacterium tuberculosis from Multiple Laboratories. J Bacteriol. 2010 July 15;192(14):3645–53.

13. Bhalla N, Nanda RK. Pangenome-wide association study reveals the selective absence of CRISPR genes (Rv2816c-19c) in drug-resistant Mycobacterium tuberculosis. Microbiol Spectr. 2024 Aug 6;12(8):e0052724.

14. Negrete-Paz AM, Vázquez-Marrufo G, Gutiérrez-Moraga A, Vázquez-Garcidueñas MaS. Pangenome Reconstruction of Mycobacterium tuberculosis as a Guide to Reveal Genomic Features Associated with Strain Clinical Phenotype. Microorganisms. 2023 June 4;11(6):1495.

15. Bespiatykh D, Bespyatykh J, Mokrousov I, Shitikov E. A Comprehensive Map of Mycobacterium tuberculosis Complex Regions of Difference. mSphere. 6(4):e00535–21.

16. He C, Cheng X, Kaisaier A, Wan J, Luo S, Ren J, et al. Effects of Mycobacterium tuberculosis lineages and regions of difference (RD) virulence gene variation on tuberculosis recurrence. Ann Transl Med. 2022 Jan;10(2):49.

17. Bahk K, Sung J, Seki M, Kim K, Kim J, Choi H, et al. Pan-lineage Mycobacterium tuberculosis reference genome for enhanced molecular diagnosis. DNA Res. 2024 Aug 1;31(4):dsae023.

18. Ofori-Anyinam B, Riley AJ, Jobarteh T, Gitteh E, Sarr B, Faal-Jawara TI, et al. Comparative genomics shows differences in the electron transport and carbon metabolic pathways of Mycobacterium africanum relative to Mycobacterium tuberculosis and suggests an adaptation to low oxygen tension. Tuberculosis. 2020 Jan 1;120:101899.

19. Bohada-Lizarazo DP, Bravo-Sanabria KD, Cárdenas-Malpica P, Rodríguez R. Comparative Genomic Analysis of Mycobacterium tuberculosis Isolates Circulating in North Santander, Colombia. Trop Med Infect Dis. 2024 Aug 28;9(9):197.

20. Tzani-Tzanopoulou P, Skliros D, Megremis S, Xepapadaki P, Andreakos E, Chanishvili N, et al. Interactions of Bacteriophages and Bacteria at the Airway Mucosa: New Insights Into the Pathophysiology of Asthma. Front Allergy. 2021 Jan 26;1:617240.

21. Morris P, Marinelli LJ, Jacobs-Sera D, Hendrix RW, Hatfull GF. Genomic Characterization of Mycobacteriophage Giles: Evidence for Phage Acquisition of Host DNA by Illegitimate Recombination. J Bacteriol. 2008 Mar;190(6):2172–82.

22. Chen J, Novick RP. Phage-mediated intergeneric transfer of toxin genes. Science. 2009 Jan 2;323(5910):139–41.

23. Gomez-Gonzalez PJ, Andreu N, Phelan JE, de Sessions PF, Glynn JR, Crampin AC, et al. An integrated whole genome analysis of Mycobacterium tuberculosis reveals insights into relationship between its genome, transcriptome and methylome. Sci Rep. 2019 Mar 26;9(1):5204.

24. Genome-wide mutational biases fuel transcriptional diversity in the Mycobacterium tuberculosis complex | Nature Communications [Internet]. [cited 2025 Sept 21]. Available from: https://www.nature.com/articles/s41467-019-11948-6

25. Pertea M, Pertea GM, Antonescu CM, Chang TC, Mendell JT, Salzberg SL. StringTie enables improved reconstruction of a transcriptome from RNA-seq reads. Nat Biotechnol. 2015 Mar;33(3):290–5.

26. Uhrig S, Ellermann J, Walther T, Burkhardt P, Fröhlich M, Hutter B, et al. Accurate and efficient detection of gene fusions from RNA sequencing data. Genome Res. 2021 Mar;31(3):448–60.

27. Nicorici D, Şatalan M, Edgren H, Kangaspeska S, Murumägi A, Kallioniemi O, et al. FusionCatcher – a tool for finding somatic fusion genes in paired-end RNA-sequencing data [Internet]. bioRxiv; 2014 [cited 2025 Sept 18]. p. 011650. Available from: https://www.biorxiv.org/content/10.1101/011650v1

28. STAR-Fusion - GDC Docs [Internet]. [cited 2025 Sept 18]. Available from: https://docs.gdc.cancer.gov/Encyclopedia/pages/STAR-Fusion/

29. rnafusion: Introduction [Internet]. [cited 2025 Sept 18]. Available from: https://nf-co.re/rnafusion/3.0.2.html

30. Ha KCH, Lalonde E, Li L, Cavallone L, Natrajan R, Lambros MB, et al. Identification of gene fusion transcripts by transcriptome sequencing in BRCA1-mutated breast cancers and cell lines. BMC Med Genomics. 2011 Oct 27;4:75.

31. Maher CA, Palanisamy N, Brenner JC, Cao X, Kalyana-Sundaram S, Luo S, et al. Chimeric transcript discovery by paired-end transcriptome sequencing. Proc Natl Acad Sci U S A. 2009 July 28;106(30):12353–8.

32. Grabherr MG, Haas BJ, Yassour M, Levin JZ, Thompson DA, Amit I, et al. Trinity: reconstructing a full-length transcriptome without a genome from RNA-Seq data. Nat Biotechnol. 2011 May 15;29(7):644–52.

33. Treangen TJ, Salzberg SL. Repetitive DNA and next-generation sequencing: computational challenges and solutions. Nat Rev Genet. 2011 Nov 29;13(1):36–46.

34. Phan V, Gao S, Tran Q, Vo NS. How genome complexity can explain the difficulty of aligning reads to genomes. BMC Bioinformatics. 2015 Dec 7;16(Suppl 17):S3.

35. Treangen TJ, Salzberg SL. Repetitive DNA and next-generation sequencing: Computational challenges and solutions. Nat Rev Genet. 2012 Jan;13(1):36–46.

36. Pelly S, Winglee K, Xia FF, Stevens RL, Bishai WR, Lamichhane G. REMap: Operon Map of M. tuberculosis. Tuberc Edinb Scotl. 2016 July;99:70–80.

37. Mahillon J, Chandler M. Insertion sequences. Microbiol Mol Biol Rev MMBR. 1998 Sept;62(3):725–74.

38. Olliver A, Vallé M, Chaslus-Dancla E, Cloeckaert A. Overexpression of the Multidrug Efflux Operon acrEF by Insertional Activation with IS1 or IS10 Elements in Salmonella enterica Serovar Typhimurium DT204 acrB Mutants Selected with Fluoroquinolones. Antimicrob Agents Chemother. 2005 Jan;49(1):289–301.

39. Williams M, Mizrahi V, Kana BD. Molybdenum cofactor: a key component of Mycobacterium tuberculosis pathogenesis? Crit Rev Microbiol. 2014 Feb;40(1):18–29.

40. Levillain F, Poquet Y, Mallet L, Mazères S, Marceau M, Brosch R, et al. Horizontal acquisition of a hypoxia-responsive molybdenum cofactor biosynthesis pathway contributed to Mycobacterium tuberculosis pathoadaptation. PLoS Pathog. 2017 Nov 27;13(11):e1006752.

41. Magalon A, Mendel RR. Biosynthesis and Insertion of the Molybdenum Cofactor. EcoSal Plus. 6(2):10.1128/ecosalplus.ESP-0006–2013.

42. De Maio F, Berisio R, Manganelli R, Delogu G. PE_PGRS proteins of Mycobacterium tuberculosis: A specialized molecular task force at the forefront of host–pathogen interaction. Virulence. 11(1):898–915.

43. Abdallah AM, Verboom T, Weerdenburg EM, Gey van Pittius NC, Mahasha PW, Jiménez C, et al. PPE and PE_PGRS proteins of Mycobacterium marinum are transported via the type VII secretion system ESX-5. Mol Microbiol. 2009 Aug;73(3):329–40.

44. Gallant J, Mouton J, Ummels R, ten Hagen-Jongman C, Kriel N, Pain A, et al. Identification of gene fusion events in Mycobacterium tuberculosis that encode chimeric proteins. NAR Genomics Bioinforma. 2020 May 18;2(2):lqaa033.

45. Chen S, Tong X, Woodard RW, D. G, Wu J, Chen J. Identification and Characterization of Bacterial Cutinase. J Biol Chem. 2008 Sept 19;283(38):25854–62.

