## Supplementary 1 (Tables and Figures) for "In-silico evidence of non-operonic fusion transcripts in *Mycobacterium tuberculosis*: algorithm optimization and signatures of genome plasticity"

Nikhil Bhalla

**Supplementary data 1**

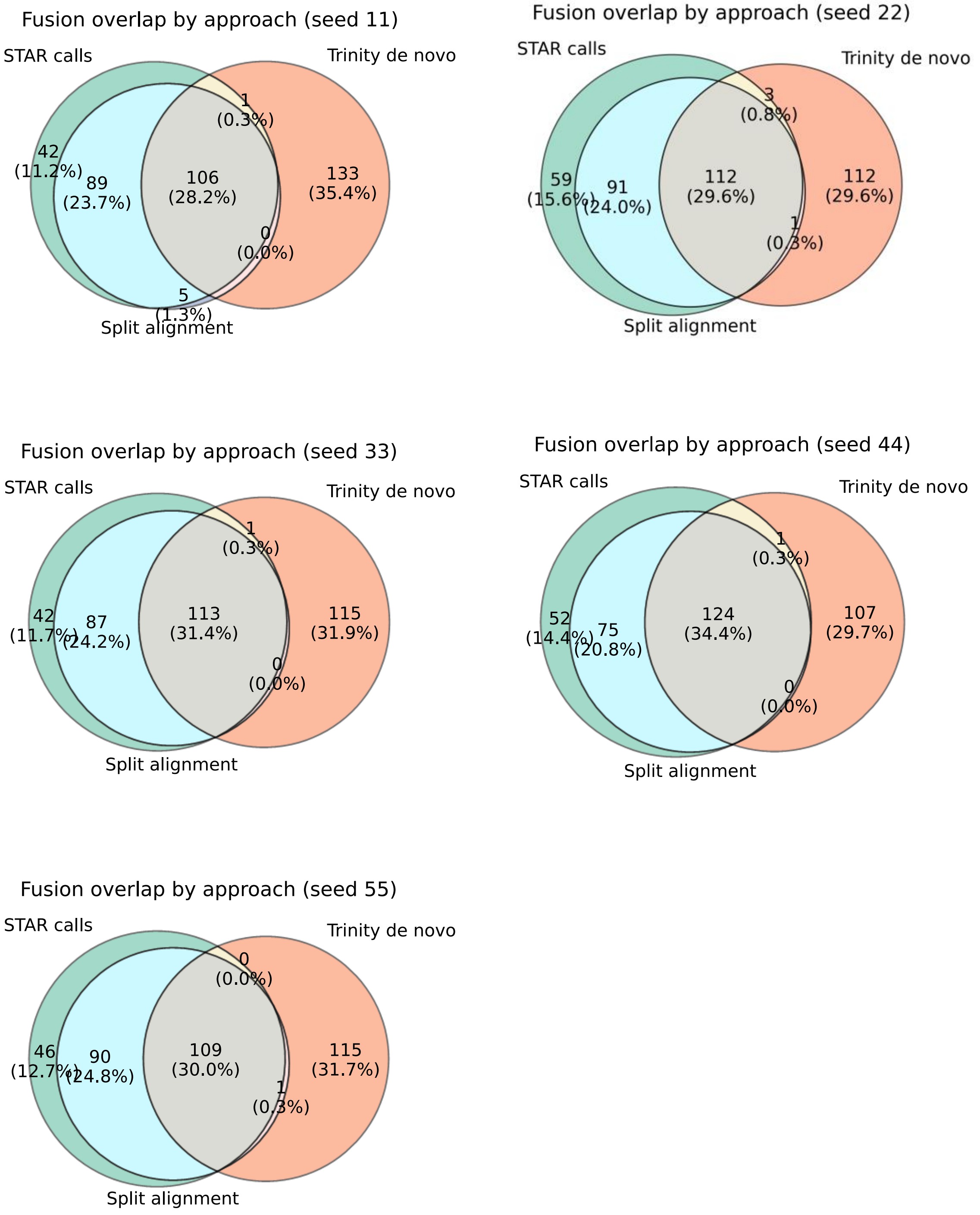


**Supplementary Figure 1:** The fusion calls made with the three approaches were intersected for all replicates (different seeds). The intersected counts were not drastically different from each other. Abbreviations: SA: split alignments, also named as split reads or approach-3 in the manuscript.

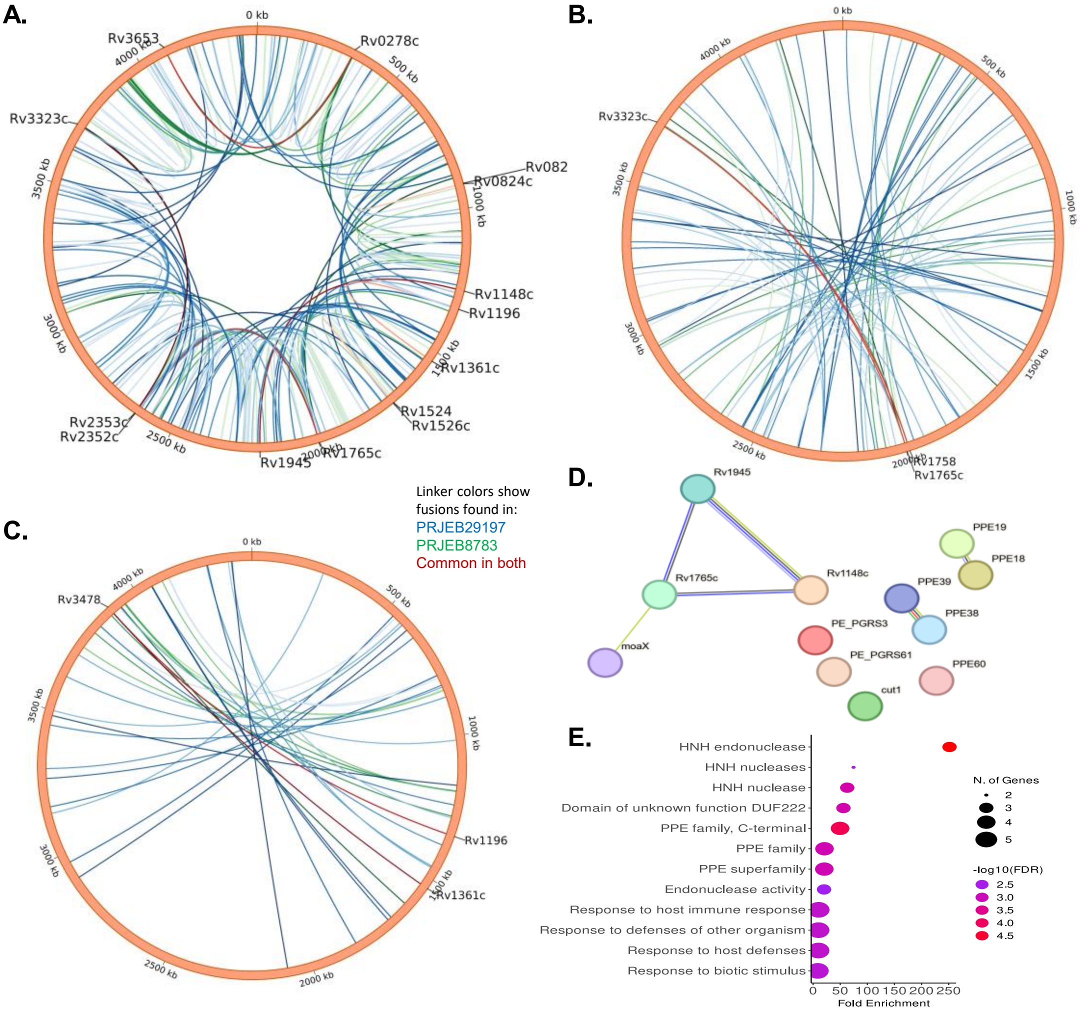


**Supplementary Figure 2: Long-distance genes form fusion transcripts.** The fusion calls from both datasets (PRJEB8783 and PRJEB29197) supported by at least 3 reads were compiled. Fusion calls from up to 6 local contiguous genes were removed. Subsequently, the fusion calls were categorized into 4 bins: first 1000 genes (bin-1), 1001-2000 genes (bin-2), 2001-3000 genes(bin-3), and 3001-last (bin-4). The fusion calls of bin-1 and bin-4 were concatenated. A: Circos representation of bin-1 and bin-4. B, C: Circos representation of bin-2 and bin-3, respectively. Only genes that formed fusion transcripts in both datasets have been labelled. D: The gene set that was found to form fusion transcripts was subjected to StringDB analysis. E: The same gene set was subjected to gene ontology clustering. Note: for visualization, the pycirclize library was used for circos diagrams. Abbreviations: kb: Kilobase, N: number, FDR: False discovery rate.

**Supplementary Table 1: Values of TP, FP, FN, precision, recall and F1 scores of the three approaches independently.** The precision =TP/(TP+FP), recall =TP/(TP+FN) and F1 scores =2.precision.recall/(precision + recall), also known as PPV, Sensitivity were determined for each approach independently. Abbreviations: TP: True positives, FP: False positives, FN: False negatives.

| | Seed | TP | FP | FN | Precision | Recall | F1 score |
|---|---|---|---|---|---|---|---|
| **Approach 1** | 11 | 194 | 202 | 6 | 0.4899 | 0.97 | 0.651 |
| | 22 | 195 | 220 | 5 | 0.4699 | 0.975 | 0.6341 |
| | 33 | 196 | 205 | 4 | 0.4888 | 0.98 | 0.6522 |
| | 44 | 194 | 207 | 6 | 0.4838 | 0.97 | 0.6456 |
| | 55 | 194 | 212 | 6 | 0.4778 | 0.97 | 0.6403 |
| **Approach 2** | 11 | 193 | 250 | 7 | 0.4357 | 0.965 | 0.6003 |
| | 22 | 194 | 287 | 6 | 0.4033 | 0.97 | 0.5698 |
| | 33 | 196 | 259 | 4 | 0.4308 | 0.98 | 0.5985 |
| | 44 | 193 | 272 | 7 | 0.4151 | 0.965 | 0.5805 |
| | 55 | 193 | 267 | 7 | 0.4196 | 0.965 | 0.5848 |
| **Apprach 3** | 11 | 111 | 303 | 89 | 0.2681 | 0.555 | 0.3616 |
| | 22 | 115 | 275 | 85 | 0.2949 | 0.575 | 0.3898 |
| | 33 | 119 | 278 | 81 | 0.2997 | 0.595 | 0.3987 |
| | 44 | 123 | 278 | 77 | 0.3067 | 0.615 | 0.4093 |
| | 55 | 109 | 285 | 91 | 0.2766 | 0.545 | 0.367 |

**Supplementary Table 2: Values of TP, FP, FN, precision, recall and F1 scores of the intersects for all differently seeded replicates.** The precision =TP/(TP+FP), recall =TP/(TP+FN) and F1 scores =2.precision.recall/(precision + recall), also known as PPV, Sensitivity were determined for each approach independently. Abbreviations: TP: True positives, FP: False positives, FN: False negatives.

| Seed | Section | TP | FP | FN | TN | Precision | Recall | F1 |
|---|---|---|---|---|---|---|---|---|
| 11 | A | 0 | 42 | 200 | 140 | 0 | 0 | 0 |
| 11 | B | 0 | 133 | 200 | 49 | 0 | 0 | 0 |
| 11 | C | 1 | 4 | 199 | 178 | 0.2 | 0.005 | 0.009756 |
| 11 | D | 0 | 1 | 200 | 181 | 0 | 0 | 0 |
| 11 | E | 87 | 2 | 113 | 180 | 0.977528 | 0.435 | 0.602076 |
| 11 | F | 0 | 0 | 200 | 182 | 0 | 0 | 0 |
| 11 | G | 106 | 0 | 94 | 182 | 1 | 0.53 | 0.69281 |
| 22 | A | 0 | 59 | 200 | 127 | 0 | 0 | 0 |
| 22 | B | 0 | 112 | 200 | 74 | 0 | 0 | 0 |
| 22 | C | 0 | 1 | 200 | 185 | 0 | 0 | 0 |
| 22 | D | 0 | 3 | 200 | 183 | 0 | 0 | 0 |
| 22 | E | 84 | 7 | 116 | 179 | 0.923077 | 0.42 | 0.57732 |
| 22 | F | 1 | 0 | 199 | 186 | 1 | 0.005 | 0.00995 |
| 22 | G | 108 | 4 | 92 | 182 | 0.964286 | 0.54 | 0.692308 |
| 33 | A | 0 | 42 | 200 | 122 | 0 | 0 | 0 |
| 33 | B | 0 | 115 | 200 | 49 | 0 | 0 | 0 |
| 33 | C | 0 | 2 | 200 | 162 | 0 | 0 | 0 |
| 33 | D | 0 | 1 | 200 | 163 | 0 | 0 | 0 |
| 33 | E | 84 | 3 | 116 | 161 | 0.965517 | 0.42 | 0.585366 |
| 33 | F | 0 | 0 | 200 | 164 | 0 | 0 | 0 |
| 33 | G | 112 | 1 | 88 | 163 | 0.99115 | 0.56 | 0.715655 |
| 44 | A | 0 | 52 | 200 | 115 | 0 | 0 | 0 |
| 44 | B | 0 | 107 | 200 | 60 | 0 | 0 | 0 |
| 44 | C | 1 | 0 | 199 | 167 | 1 | 0.005 | 0.00995 |
| 44 | D | 0 | 1 | 200 | 166 | 0 | 0 | 0 |
| 44 | E | 70 | 5 | 130 | 162 | 0.933333 | 0.35 | 0.509091 |
| 44 | F | 0 | 0 | 200 | 167 | 0 | 0 | 0 |
| 44 | G | 122 | 2 | 78 | 165 | 0.983871 | 0.61 | 0.753086 |
| 55 | A | 0 | 46 | 200 | 125 | 0 | 0 | 0 |
| 55 | B | 0 | 115 | 200 | 56 | 0 | 0 | 0 |
| 55 | C | 1 | 1 | 199 | 170 | 0.5 | 0.005 | 0.009901 |
| 55 | D | 0 | 0 | 200 | 171 | 0 | 0 | 0 |
| 55 | E | 85 | 5 | 115 | 166 | 0.944444 | 0.425 | 0.586207 |
| 55 | F | 0 | 1 | 200 | 170 | 0 | 0 | 0 |
| 55 | A | 106 | 3 | 94 | 168 | 0.972477 | 0.53 | 0.686084 |
| 11 | B | 193 | 2 | 7 | 0 | 0.989744 | 0.965 | 0.977215 |
| 22 | C | 192 | 11 | 8 | 0 | 0.945813 | 0.96 | 0.952854 |
| 33 | D | 196 | 4 | 4 | 0 | 0.98 | 0.98 | 0.98 |
| 44 | E | 192 | 7 | 8 | 0 | 0.964824 | 0.96 | 0.962406 |
| 55 | F | 191 | 8 | 9 | 0 | 0.959799 | 0.955 | 0.957393 |

**Supplementary Table 3: Gene ontology details of genes forming fusion transcripts in Mtb**
The gene set was analysed using ShinyGO online tool.

| Enrichment FDR | nGenes | Pathway Genes | Fold Enrichment | Pathway | Genes |
|---|---|---|---|---|---|
| 2.65E-07 | 9 | 50 | 18.58154 | PE family | RV0278C RV0279C RV0747 RV0980C RV1068C RV1452C RV1759C RV2615C RV3511 |
| 1.40E-06 | 10 | 88 | 11.73077 | PE-PGRS family, N-terminal | RV0278C RV0279C RV0747 RV0980C RV1068C RV1452C RV1759C RV1840C RV2615C RV3511 |
| 0.000557 | 3 | 4 | 77.42308 | HNH endonuclease | RV1148C RV1765C RV1945 |
| 0.000618 | 3 | 5 | 61.93846 | Transposase, mutator type, and Transposase activity | RV1199C RV2512C RV3115 |
| 0.000618 | 5 | 34 | 15.181 | ER to Golgi transport vesicle membrane | RV1148C RV1199C RV1765C RV2512C RV3115 |
| 0.000618 | 5 | 34 | 15.181 | Membrane coat | RV1148C RV1199C RV1765C RV2512C RV3115 |
| 0.000618 | 5 | 34 | 15.181 | Vesicle coat | RV1148C RV1199C RV1765C RV2512C RV3115 |
| 0.000618 | 5 | 34 | 15.181 | COPII vesicle coat | RV1148C RV1199C RV1765C RV2512C RV3115 |
| 0.000618 | 5 | 34 | 15.181 | Transport vesicle | RV1148C RV1199C RV1765C RV2512C RV3115 |
| 0.000618 | 5 | 34 | 15.181 | COPII-coated ER to Golgi transport vesicle | RV1148C RV1199C RV1765C RV2512C RV3115 |
| 0.000618 | 5 | 34 | 15.181 | Transport vesicle membrane | RV1148C RV1199C RV1765C RV2512C RV3115 |
| 0.000618 | 5 | 34 | 15.181 | Coated membrane | RV1148C RV1199C RV1765C RV2512C RV3115 |
| 0.000711 | 5 | 37 | 13.9501 | Golgi-associated vesicle | RV1148C RV1199C RV1765C RV2512C RV3115 |
| 0.000711 | 5 | 37 | 13.9501 | Coated vesicle | RV1148C RV1199C RV1765C RV2512C RV3115 |
| 0.000711 | 5 | 37 | 13.9501 | Golgi-associated vesicle membrane | RV1148C RV1199C RV1765C RV2512C RV3115 |
| 0.000711 | 5 | 37 | 13.9501 | Coated vesicle membrane | RV1148C RV1199C RV1765C RV2512C RV3115 |
| 0.001417 | 5 | 43 | 12.00358 | Cytoplasmic vesicle membrane | RV1148C RV1199C RV1765C RV2512C RV3115 |
| 0.001499 | 5 | 44 | 11.73077 | Vesicle membrane | RV1148C RV1199C RV1765C RV2512C RV3115 |
| 0.00209 | 6 | 77 | 8.043956 | Organelle membrane | RV1148C RV1199C RV1371 RV1765C RV2512C RV3115 |
| 0.003054 | 7 | 120 | 6.021795 | Membrane protein complex | RV1148C RV1199C RV1765C RV1945 RV1964 RV2512C RV3115 |
| 0.00313 | 3 | 11 | 28.15385 | Mixed, incl. Decaprenyl diphosphate synthase-like, and Cell wall | RV3511 RV3512 RV3653 |
| 0.003328 | 5 | 54 | 9.558405 | Bounding membrane of organelle | RV1148C RV1199C RV1765C RV2512C RV3115 |
| 0.003955 | 5 | 57 | 9.055331 | Cytoplasmic vesicle | RV1148C RV1199C RV1765C RV2512C RV3115 |
| 0.003955 | 5 | 57 | 9.055331 | Intracellular vesicle | RV1148C RV1199C RV1765C RV2512C RV3115 |
| 0.007059 | 3 | 15 | 20.64615 | Mostly uncharacterized, incl. DDE transposase retroviral integrase sub-family, and Transposase, IS111A/IS1328/IS1533, N-terminal | RV0395 RV2808 RV3327 |
| 0.007303 | 3 | 16 | 19.35577 | Mixed, incl. WD40/YVTN repeat-like-containing domain superfamily, and Polyketide cyclase/dehydrase | RV0980C RV1059 RV2615C |
| 0.007303 | 4 | 38 | 10.8664 | Transposase activity | RV1199C RV2512C RV3115 RV3327 |
| 0.007303 | 5 | 66 | 7.820513 | Endomembrane system | RV1148C RV1199C RV1765C RV2512C RV3115 |
| 0.007303 | 5 | 68 | 7.590498 | Vesicle | RV1148C RV1199C RV1765C RV2512C RV3115 |
| 0.007303 | 3 | 16 | 19.35577 | HNH nuclease | RV1148C RV1765C RV1945 |
| 0.00782 | 4 | 39 | 10.58777 | Transposition, DNA-mediated | RV1199C RV2512C RV3115 RV3327 |
| 0.008469 | 2 | 4 | 51.61538 | Carbon fixation | RV3280 RV3281 |
| 0.0094 | 3 | 18 | 17.20513 | Domain of unknown function DUF222 | RV1148C RV1765C RV1945 |
| 0.01247 | 2 | 5 | 41.29231 | Carbon fixation, and Thiosulphate sulfurtransferase, conserved site | RV3280 RV3281 |

| | | | | | |
|---|---|---|---|---|---|
| 0.01247 | 2 | 5 | 41.29231 | Mixed, incl. Transposase, IS111A/IS1328/IS1533, N-terminal, and Glycoside hydrolase superfamily | RV2808 RV3327 |
| 0.01247 | 2 | 5 | 41.29231 | propionyl-CoA carboxylase activity | RV3280 RV3281 |
| 0.016732 | 2 | 6 | 34.41026 | Mixed, incl. PIN domain toxin, and Ribbon-helix-helix protein, CopG | RV0278C RV0747 |
| 0.016732 | 2 | 6 | 34.41026 | acetyl-CoA carboxylase activity | RV3280 RV3281 |
| 0.016732 | 2 | 6 | 34.41026 | acetyl-CoA carboxylase complex | RV3280 RV3281 |
| 0.016732 | 2 | 6 | 34.41026 | CoA carboxylase activity | RV3280 RV3281 |
| 0.020347 | 3 | 25 | 12.38769 | Mixed, incl. Transposase, mutator type, and Transposase activity | RV1199C RV2512C RV3115 |
| 0.020817 | 3 | 26 | 11.91124 | Mostly uncharacterized, incl. Methyltransferase domain, and DegT/DnrJ/EryC1/StrS aminotransferase | RV0395 RV2808 RV3327 |
| 0.020817 | 2 | 7 | 29.49451 | Mixed, incl. Glucose-methanol-choline oxidoreductase, and tRNA threonylcarbamoyladenosine dehydratase | RV1758 RV2338C |
| 0.020817 | 2 | 7 | 29.49451 | Ligase activity, forming carbon-carbon bonds | RV3280 RV3281 |
| 0.022757 | 3 | 27 | 11.47009 | Mostly uncharacterized, incl. Metalloprotease TldD/PmbA superfamily, and ESAT-6-like superfamily | RV3511 RV3512 RV3653 |
| 0.029813 | 4 | 62 | 6.66005 | Transposition | RV1199C RV2512C RV3115 RV3327 |
| 0.032853 | 2 | 9 | 22.94017 | HNH nucleases | RV1765C RV1945 |
| 0.039983 | 2 | 10 | 20.64615 | Mixed, incl. WD40/YVTN repeat-like-containing domain superfamily, and Dihydrodipicolinate reductase, N-terminal | RV0980C RV1059 |
| 0.052143 | 4 | 74 | 5.580042 | Repeat | RV0278C RV0747 RV0980C RV1759C |
| 0.054587 | 2 | 12 | 17.20513 | Mostly uncharacterized, incl. Vancomycin metabolic process, and Domain of unknown function DUF4185 | RV1759C RV1765C |
| 0.083657 | 2 | 15 | 13.7641 | Mixed, incl. Cutinase/acetylxylan esterase, and Carboxylesterase, type B | RV1758 RV2338C |
| 0.089483 | 7 | 253 | 2.856187 | Extracellular region | RV0278C RV0279C RV0747 RV1758 RV1759C RV3327 RV3512 |
| 0.109398 | 3 | 50 | 6.193846 | Endonuclease activity | RV1148C RV1765C RV1945 |
| 0.111291 | 2 | 18 | 11.47009 | Domain of unknown function DUF222, and DNA methylation or demethylation | RV1148C RV1945 |
| 0.111291 | 2 | 18 | 11.47009 | Mixed, incl. PE family, and PIN domain toxin | RV0278C RV0747 |
| 0.113471 | 3 | 52 | 5.955621 | Mixed, incl. PE-PGRS family, N-terminal, and PPE family | RV3511 RV3512 RV3653 |
| 0.113471 | 4 | 96 | 4.301282 | DNA recombination | RV1199C RV2512C RV3115 RV3327 |
| 0.117447 | 2 | 19 | 10.8664 | CoA carboxylase activity, and Fatty acid biosynthesis | RV3280 RV3281 |
| 0.175205 | 2 | 24 | 8.602564 | Mostly uncharacterized, incl. Domain of unknown function DUF222, and Viral process | RV1759C RV1765C |
| 0.175205 | 4 | 112 | 3.686813 | Catalytic activity, acting on DNA | RV1199C RV2512C RV3115 RV3327 |

| | | | | | |
|---|---|---|---|---|---|
| 0.183388 | 2 | 25 | 8.258462 | Mixed, incl. CoA carboxylase activity, and Fatty acid biosynthesis | RV3280 RV3281 |
| 0.183388 | 2 | 25 | 8.258462 | Domain of unknown function DUF222, and Rhodanese Homology Domain | RV1148C RV1945 |
| 0.192562 | 32 | 2681 | 1.232146 | Cellular anatomical entity | RV0278C RV0279C RV0747 RV0980C RV1068C RV1148C RV1199C RV1361C RV1371 RV1452C RV1758 RV1759C RV1765C RV1840C RV1945 RV1964 RV1977 RV2277C RV2282C RV2338C RV2353C RV2512C RV3115 RV3280 RV3281 RV3323C RV3327 RV3478 RV3511 RV3512 RV3653 RV3734C |
| 0.197175 | 8 | 382 | 2.161901 | Nucleic acid metabolic process | RV1148C RV1199C RV1765C RV1945 RV2338C RV2512C RV3115 RV3327 |
| 0.197175 | 1 | 3 | 34.41026 | Peptidase M48 | RV1977 |
| 0.197175 | 1 | 3 | 34.41026 | Molybdopterin biosynthesis MoaE | RV3323C |
| 0.197175 | 2 | 27 | 7.646724 | PPE family, C-terminal | RV1361C RV3478 |
| 0.197175 | 1 | 3 | 34.41026 | Glycerophosphodiester phosphodiesterase domain | RV2277C |
| 0.197175 | 1 | 3 | 34.41026 | Molybdopterin biosynthesis MoaE subunit superfamily | RV3323C |
| 0.197175 | 1 | 3 | 34.41026 | NHL repeat | RV0980C |
