## Supplementary 1 (Codes) for "In-silico evidence of non-operonic fusion transcripts in *Mycobacterium tuberculosis*: algorithm optimization and signatures of genome plasticity"

Nikhil Bhalla

**Supplementary-2 (contains scripts used in the study)**

### 2.1. Fusion simulation CDS preparation script

```
#!/usr/bin/env python3
import argparse
import random
import sys
from collections import namedtuple
from pathlib import Path
from Bio import SeqIO
from Bio.Seq import Seq
def parse_args():
    parser = argparse.ArgumentParser()
    parser.add_argument("-i", "--cds_fasta", required=True)
    parser.add_argument("-n", "--num_fusions", type=int, default=200)
    parser.add_argument("--prop_inframe", type=float, default=0.6)
    parser.add_argument("--min_len_nt", type=int, default=300)
    parser.add_argument("--min_head_nt", type=int, default=150)
    parser.add_argument("--min_tail_nt", type=int, default=150)
    parser.add_argument("--require_no_stop", action="store_true")
    parser.add_argument("--seed", type=int, default=42)
    parser.add_argument("-o", "--out_fasta", default="fusions.fa")
    parser.add_argument("-t", "--out_truth", default="fusions.tsv")
    return parser.parse_args()
def has_internal_stop(nt_seq):
    return "*" in str(Seq(str(nt_seq)).translate(to_stop=False))[:-1]
def main():
    args = parse_args()
    random.seed(args.seed)
    records = [r for r in SeqIO.parse(args.cds_fasta, "fasta") if len(r.seq) >= args.min_len_nt]
    if len(records) < 2:
        sys.exit("Not enough CDS records of sufficient length.")
    Pool = namedtuple("PoolRec", "id seq")
    pool = [Pool(r.id, Seq(str(r.seq).upper().replace("U", "T"))) for r in records]
    fasta_lines = []
    truth_lines = ["fusion_id\tcdsA_id\tcutA_nt\tcdsB_id\tcutB_nt\tframe_status\tlenA\tlenB\n"]
    made = 0
    tries = 0
    max_tries = args.num_fusions * 20
    while made < args.num_fusions and tries < max_tries:
        tries += 1
        A, B = random.sample(pool, 2)
        lenA, lenB = len(A.seq), len(B.seq)
        if random.random() < args.prop_inframe:
            validA = [p for p in range(args.min_head_nt, lenA - args.min_head_nt) if p % 3 == 0]
            validB = [p for p in range(args.min_tail_nt, lenB - args.min_tail_nt) if p % 3 == 0]
            if not validA or not validB:
                continue
            cutA = random.choice(validA)
            cutB = random.choice(validB)
            frame = "inframe"
        else:
            cutA = random.randrange(args.min_head_nt, lenA - args.min_head_nt)
            cutB = random.randrange(args.min_tail_nt, lenB - args.min_tail_nt)
            if (cutA % 3) == (cutB % 3):
                cutB = cutB + 1 if (cutB + 1) < (lenB - args.min_tail_nt) else max(args.min_tail_nt,
cutB - 1)
            frame = "outframe"
        head = A.seq[:cutA]
        tail = B.seq[cutB:]
        if len(head) < args.min_head_nt or len(tail) < args.min_tail_nt:
            continue
        fusion_seq = head + tail
        if args.require_no_stop and has_internal_stop(fusion_seq):
            continue
        fid = f"FUS_{made:04d}|{A.id}:{cutA}|{B.id}:{cutB}|{frame}"
        fasta_lines.append(f">{fid}\n{fusion_seq}\n")
        truth_lines.append("\t".join(map(str, [fid, A.id, cutA, B.id, cutB, frame, lenA, lenB])) +
"\n")
        made += 1
    if made < args.num_fusions:
        sys.stderr.write(f"Warning: generated {made}/{args.num_fusions} fusions\n")
    Path(args.out_fasta).parent.mkdir(parents=True, exist_ok=True)
    with open(args.out_fasta, "w") as out_fa:
```

```
        out_fa.writelines(fasta_lines)
    with open(args.out_truth, "w") as out_tsv:
        out_tsv.writelines(truth_lines)
if __name__ == "__main__":
    main()
```

### 2.2. Approach 1

```
#!/usr/bin/env bash
set -euo pipefail
minimap2 -t 16 -ax sr reference.fasta reads_1.fq reads_2.fq | samtools sort -o mapped.bam
samtools index mapped.bam
awk -F'\t' 'BEGIN{OFS="\t"} $3=="CDS"{
  p=""
  split($9,a,";")
  for(i in a){gsub(/^ +| +$/, "", a[i]); if(a[i]~"^Parent=") p=substr(a[i],8)}
  if(p!="") print $1,$4-1,$5,p,".",$7
}' annotations.gff3 > cds.bed
python3 splitmap2csv.py -b mapped.bam --cds_bed cds.bed -w 10 --min_support 3 -o calls.csv
python3 score_pairs.py --calls_csv calls.csv --truth_tsv truth_fusions.tsv --out_prefix
results
```

### 2.3. Approach 2

```
#!/usr/bin/env bash
set -euo pipefail
awk 'BEGIN{OFS="\t"} $3=="CDS"{
  gene=""
  split($9,a,";")
  for(i in a){gsub(/^ +| +$/, "", a[i]); if(a[i]~/^Parent=/) gene=substr(a[i],8)}
  if(gene!="") print $1,"sim","exon",$4,$5,".",$7,".","gene_id \""gene"\"; transcript_id \""gene"\";
gene_name \""gene"\";"
}' annotations.gff3 > genes.exon.gtf
STAR --runThreadN 12 --runMode genomeGenerate --genomeDir star_idx --genomeFastaFiles genome.fasta --
sjdbGTFfile genes.exon.gtf
STAR --runThreadN 12 --genomeDir star_idx --readFilesIn reads_1.fq reads_2.fq --outSAMtype BAM
SortedByCoordinate --alignIntronMax 1 --alignSJoverhangMin 999 --alignSJDBoverhangMin 999 --
chimOutType Junctions WithinBAM HardClip --chimSegmentMin 10 --chimJunctionOverhangMin 10 --
chimMultimapNmax 50 --outFileNamePrefix star_
tail -n +2 star_Chimeric.out.junction | awk 'BEGIN{OFS="\t"} ($2~/^[0-9]+$/ && $5~/^[0-9]+$/){print
$1,$2,$3,$4,$5,$6}' | sort -k1,1 -k2,2n -k3,3 -k4,4 -k5,5n -k6,6 | uniq -c | awk
'BEGIN{OFS="\t"}{print $2,$3,$4,$5,$6,$7,$1}' > star_junctions.counts.tsv
awk 'BEGIN{OFS="\t"}{key=$1"|"$2"|"$3"|"$4"|"$5"|"$6"|"$7;s=$2-3;if(s<0)s=0;e=$2+2;print $1,s,e,key}'
star_junctions.counts.tsv > sideA.slop.bed
awk 'BEGIN{OFS="\t"}{key=$1"|"$2"|"$3"|"$4"|"$5"|"$6"|"$7;s=$5-3;if(s<0)s=0;e=$5+2;print $4,s,e,key}'
star_junctions.counts.tsv > sideB.slop.bed
bedtools intersect -wa -wb -a sideA.slop.bed -b cds.bed > A2cds.tsv
bedtools intersect -wa -wb -a sideB.slop.bed -b cds.bed > B2cds.tsv
bedtools intersect -u -a sideA.slop.bed -b mask.bed 2>/dev/null | cut -f4 > mask.keysA || true
bedtools intersect -u -a sideB.slop.bed -b mask.bed 2>/dev/null | cut -f4 > mask.keysB || true
cat mask.keysA mask.keysB 2>/dev/null | sort -u > mask.keys || true
python3 - <<'PY'
import csv
def parse_key(text):
    cA,pA,stA,cB,pB,stB,sup = text.split("|")
    return cA,int(pA),stA,cB,int(pB),stB,int(sup)
def load_hits(path):
    hits = {}
    try:
        with open(path) as handle:
            for line in handle:
                parts = line.rstrip("\n").split("\t")
                if len(parts) >= 8:
                    hits.setdefault(parts[3], set()).add(parts[7])
    except FileNotFoundError:
        pass
    return hits
masked = set()
try:
```

```
    with open("mask.keys") as handle:
        masked = {line.strip() for line in handle}
except FileNotFoundError:
    pass
geneA = load_hits("A2cds.tsv")
geneB = load_hits("B2cds.tsv")
with open("star_calls.csv", "w", newline="") as handle:
    writer = csv.writer(handle)
writer.writerow(["approach","geneA","geneB","chrA","posA","strandA","chrB","posB","strandB","support_s
hortreads","support_longreads","in_frame","orf_len","masked_flag","notes"])
    for key in sorted(geneA.keys() & geneB.keys()):
        cA,pA,stA,cB,pB,stB,sup = parse_key(key)
        left = ";".join(sorted(geneA[key]))
        right = ";".join(sorted(geneB[key]))
        if left and right:
            writer.writerow(["star", left, right, cA, pA, stA, cB, pB, stB, sup, 0, "", "", "1" if key
in masked else "0", ""])
PY
python3 score_pairs.py --calls_csv star_calls.csv --truth
```

### 2.4. Approach – 3

```
#!/usr/bin/env bash
set -euo pipefail
Trinity --seqType fq --left reads_1.fq --right reads_2.fq --CPU 16 --max_memory 40G --output
trinity_out --full_cleanup
threads=8
minimap2 -t "$threads" -ax asm5 genome.fasta trinity_out/Trinity.fasta | samtools sort -@ "$threads" -
o trinity.bam
samtools index trinity.bam
python3 - <<'PY'
import os
import csv
import tempfile
import subprocess
from collections import defaultdict
import pysam
bam_path = "trinity.bam"
cds_bed = "cds.bed"
mask_bed = "mask.bed" if os.path.exists("mask.bed") and os.path.getsize("mask.bed") else None
window = 15
min_support = 1
slop = 10
def bed_intersect(a, b, args=("-wa", "-wb")):
    out = subprocess.check_output(["bedtools", "intersect", *args, "-a", a, "-b", b], text=True)
    return [line for line in out.strip().splitlines() if line]
def cigar_ref_len(items):
    return sum(length for op, length in (items or []) if op in (0, 2, 3, 7, 8))
def parse_sa(text):
    out = []
    for token in text.split(";"):
        if not token:
            continue
        fields = token.split(",")
        if len(fields) < 4:
            continue
        try:
            out.append((fields[0], int(fields[1]), fields[2], fields[3]))
        except ValueError:
            pass
    return out
bam = pysam.AlignmentFile(bam_path, "rb")
clusters = {}
for read in bam.fetch(until_eof=True):
    if read.is_unmapped or not read.has_tag("SA"):
        continue
    chr_a = read.reference_name
    strand_a = "-" if read.is_reverse else "+"
    start_a = read.reference_start + 1
    end_a = start_a + cigar_ref_len(read.cigartuples) - 1
    pos_a = end_a
    for chr_b, pos_b, strand_b, _ in parse_sa(read.get_tag("SA")):
        bin_a = round(pos_a / window)
```

```
        bin_b = round(pos_b / window)
        key = (chr_a, strand_a, chr_b, strand_b, bin_a, bin_b)
        clusters.setdefault(key, [0, 0, 0])
        clusters[key][0] += pos_a
        clusters[key][1] += pos_b
        clusters[key][2] += 1
bam.close()
junctions = []
for (chr_a, strand_a, chr_b, strand_b, _, _), (sum_a, sum_b, count) in clusters.items():
    if count < min_support:
        continue
    pos_a = round(sum_a / count)
    pos_b = round(sum_b / count)
    junction_id = f"{chr_a}:{pos_a}:{strand_a}|{chr_b}:{pos_b}:{strand_b}"
    junctions.append((junction_id, chr_a, pos_a, strand_a, chr_b, pos_b, strand_b, count))
tmp_dir = tempfile.mkdtemp(prefix="denovo_")
side_a = os.path.join(tmp_dir, "A.bed")
side_b = os.path.join(tmp_dir, "B.bed")
with open(side_a, "w") as fh_a, open(side_b, "w") as fh_b:
    for junction_id, chr_a, pos_a, strand_a, chr_b, pos_b, strand_b, count in junctions:
        fh_a.write(f"{chr_a}\t{max(0, pos_a - 1 - slop)}\t{pos_a + slop}\t{junction_id}\n")
        fh_b.write(f"{chr_b}\t{max(0, pos_b - 1 - slop)}\t{pos_b + slop}\t{junction_id}\n")
a_hits = bed_intersect(side_a, cds_bed) if os.path.getsize(side_a) else []
b_hits = bed_intersect(side_b, cds_bed) if os.path.getsize(side_b) else []
left_genes = defaultdict(set)
right_genes = defaultdict(set)
for line in a_hits:
    parts = line.split("\t")
    if len(parts) >= 8:
        left_genes[parts[3]].add(parts[7])
for line in b_hits:
    parts = line.split("\t")
    if len(parts) >= 8:
        right_genes[parts[3]].add(parts[7])
masked = defaultdict(lambda: "0")
if mask_bed and os.path.getsize(side_a) and os.path.getsize(side_b):
    for bed_path in (side_a, side_b):
        hits = bed_intersect(bed_path, mask_bed, args=("-u",))
        for line in hits:
            masked[line.split("\t")[3]] = "1"
with open("denovo_calls.csv", "w", newline="") as handle:
    writer = csv.writer(handle)
    writer.writerow(["approach", "geneA", "geneB", "chrA", "posA", "strandA", "chrB", "posB",
"strandB", "support_shortreads", "support_longreads", "in_frame", "orf_len", "masked_flag", "notes"])
    for junction_id, chr_a, pos_a, strand_a, chr_b, pos_b, strand_b, count in junctions:
        genes_left = ";".join(sorted(left_genes.get(junction_id, [])))
        genes_right = ";".join(sorted(right_genes.get(junction_id, [])))
        if genes_left and genes_right:
            writer.writerow(["C_denovo", genes_left, genes_right, chr_a, pos_a, strand_a, chr_b,
pos_b, strand_b, count, 0, "", "", masked[junction_id], "contigs"])
PY
```

### 2.5. Script convert SA map coordinates into junctions (used in approach-1).

```
#!/usr/bin/env python3
import argparse
import csv
import os
import subprocess
import sys
import tempfile
from collections import defaultdict
try:
    import pysam
except ImportError:
    sys.stderr.write("pysam required\n")
    sys.exit(1)
def bed_intersect(a_path, b_path, args=("-wa", "-wb")):
    cmd = ["bedtools", "intersect", *args, "-a", a_path, "-b", b_path]
    out = subprocess.check_output(cmd, text=True)
    return [line for line in out.strip().splitlines() if line]
```

```
def cigar_ref_len(cigar):
    return sum(count for op, count in (cigar or []) if op in (0, 2, 3, 7, 8))
def parse_sa(sa_tag):
    out = []
    for token in sa_tag.split(";"):
        if not token:
            continue
        fields = token.split(",")
        if len(fields) < 4:
            continue
        try:
            out.append((fields[0], int(fields[1]), fields[2], fields[3]))
        except ValueError:
            pass
    return out
def write_csv_header(handle):
csv.writer(handle).writerow(["approach","geneA","geneB","chrA","posA","strandA","chrB","posB","strandB
","support_shortreads","support_longreads","in_frame","orf_len","masked_flag","notes"])
def cluster_junctions(bam_handle, window, max_sa):
    clusters = {}
    reads_with_sa = 0
    sa_segments_used = 0
    for alignment in bam_handle.fetch(until_eof=True):
        if alignment.is_unmapped or not alignment.has_tag("SA"):
            continue
        reads_with_sa += 1
        chr_a = alignment.reference_name
        strand_a = "-" if alignment.is_reverse else "+"
        start_a = alignment.reference_start + 1
        end_a = start_a + cigar_ref_len(alignment.cigartuples) - 1
        pos_a = end_a
        for chr_b, pos_b, strand_b, _ in parse_sa(alignment.get_tag("SA"))[:max_sa]:
            sa_segments_used += 1
            bin_a = round(pos_a / window)
            bin_b = round(pos_b / window)
            key = (chr_a, strand_a, chr_b, strand_b, bin_a, bin_b)
            clusters.setdefault(key, [0, 0, 0])
            clusters[key][0] += pos_a
            clusters[key][1] += pos_b
            clusters[key][2] += 1
    return clusters, reads_with_sa, sa_segments_used
def materialize_clusters(clusters, min_support):
    junctions = []
    for (chr_a, strand_a, chr_b, strand_b, _, _), (sum_a, sum_b, count) in clusters.items():
        if count < min_support:
            continue
        pos_a = round(sum_a / count)
        pos_b = round(sum_b / count)
        junction_id = f"{chr_a}:{pos_a}:{strand_a}|{chr_b}:{pos_b}:{strand_b}"
        junctions.append((junction_id, chr_a, pos_a, strand_a, chr_b, pos_b, strand_b, count))
    return junctions
def annotate_junctions(junctions, cds_bed, mask_bed):
    tmpdir = tempfile.mkdtemp(prefix="splitmap2csv_")
    side_a_bed = os.path.join(tmpdir, "A.bed")
    side_b_bed = os.path.join(tmpdir, "B.bed")
    with open(side_a_bed, "w") as handle_a, open(side_b_bed, "w") as handle_b:
        for junction_id, chr_a, pos_a, strand_a, chr_b, pos_b, strand_b, _ in junctions:
            handle_a.write(f"{chr_a}\t{pos_a - 1}\t{pos_a}\t{junction_id}\n")
            handle_b.write(f"{chr_b}\t{pos_b - 1}\t{pos_b}\t{junction_id}\n")
    annotations_a = bed_intersect(side_a_bed, cds_bed)
    annotations_b = bed_intersect(side_b_bed, cds_bed)
    genes_a = defaultdict(set)
    genes_b = defaultdict(set)
    for line in annotations_a:
        parts = line.split("\t")
        if len(parts) >= 8:
            genes_a[parts[3]].add(parts[7])
    for line in annotations_b:
        parts = line.split("\t")
        if len(parts) >= 8:
            genes_b[parts[3]].add(parts[7])
    masked = defaultdict(lambda: "0")
    if mask_bed and os.path.exists(mask_bed) and os.path.getsize(mask_bed) > 0:
```

```
        for bed_path in (side_a_bed, side_b_bed):
            hits = bed_intersect(bed_path, mask_bed, args=("-u",))
            for line in hits:
                masked[line.split("\t")[3]] = "1"
    return genes_a, genes_b, masked
def main():
    parser = argparse.ArgumentParser()
    parser.add_argument("-b", "--bam", required=True)
    parser.add_argument("--cds_bed", required=True)
    parser.add_argument("--mask_bed", default=None)
    parser.add_argument("-w", "--window", type=int, default=10)
    parser.add_argument("-o", "--out_csv", default="A_splitmap_calls.csv")
    parser.add_argument("--min_support", type=int, default=1)
    parser.add_argument("--max_sa", type=int, default=4)
    args = parser.parse_args()
    window = max(1, args.window)
    bam_handle = pysam.AlignmentFile(args.bam, "rb")
    clusters, reads_with_sa, sa_segments_used = cluster_junctions(bam_handle, window, args.max_sa)
    bam_handle.close()
    if not clusters:
        sys.stderr.write("No chimeric SA alignments found.\n")
        with open(args.out_csv, "w", newline="") as handle:
            write_csv_header(handle)
        return
    junctions = materialize_clusters(clusters, args.min_support)
    if not junctions:
        sys.stderr.write("No junction clusters passed min_support.\n")
    genes_a, genes_b, masked = annotate_junctions(junctions, args.cds_bed, args.mask_bed)
    with open(args.out_csv, "w", newline="") as handle:
        write_csv_header(handle)
        writer = csv.writer(handle)
        for junction_id, chr_a, pos_a, strand_a, chr_b, pos_b, strand_b, support in junctions:
            genes_left = ";".join(sorted(genes_a.get(junction_id, [])))
            genes_right = ";".join(sorted(genes_b.get(junction_id, [])))
            if not genes_left or not genes_right:
                continue
            writer.writerow(["A_splitmap", genes_left, genes_right, chr_a, pos_a, strand_a, chr_b,
pos_b, strand_b, support, 0, "", "", masked[junction_id], ""])
if __name__ == "__main__":
    main()
```

### 2.6. Scoring script

```
#!/usr/bin/env python3
import argparse
import os
import pandas as pd
def normalize_id(value: str) -> str:
    if value is None:
        return ""
    text = str(value)
    for key in ("ID=", "cds_id=", "transcript_id=", "gene="):
        idx = text.find(key)
        if idx != -1:
            rem = text[idx + len(key) :]
            for sep in (";", " ", ","):
                cut = rem.find(sep)
                if cut != -1:
                    rem = rem[:cut]
                    break
            return rem
    return text.strip()
def expand_rows(row):
    genes_a = [g for g in row["geneA_norm"].split(";") if g]
    genes_b = [g for g in row["geneB_norm"].split(";") if g]
    out = []
    for gene_a in genes_a:
        for gene_b in genes_b:
            out.append({"geneA": gene_a, "geneB": gene_b, "chrA": row["chrA"], "posA": row["posA"],
"strandA": row["strandA"], "chrB": row["chrB"], "posB": row["posB"], "strandB": row["strandB"],
"support_shortreads": row["support_shortreads"], "masked_flag": row.get("masked_flag", 0), "approach":
row["approach"]})
    return out
```

```
def compute_metrics(tp_pairs, fp_pairs, fn_pairs):
    tp = len(tp_pairs)
    fp = len(fp_pairs)
    fn = len(fn_pairs)
    precision = tp / (tp + fp) if (tp + fp) else 0.0
    recall = tp / (tp + fn) if (tp + fn) else 0.0
    f1 = 2 * precision * recall / (precision + recall) if (precision + recall) else 0.0
    return tp, fp, fn, precision, recall, f1
def ensure_parent_dir(path_prefix: str) -> None:
    parent = os.path.dirname(path_prefix) or "."
    os.makedirs(parent, exist_ok=True)
def main():
    parser = argparse.ArgumentParser()
    parser.add_argument("--calls_csv", required=True)
    parser.add_argument("--truth_tsv", required=True)
    parser.add_argument("--out_prefix", default="work/score_splitmap")
    parser.add_argument("--unordered", action="store_true")
    args = parser.parse_args()
    calls = pd.read_csv(args.calls_csv)
    calls["geneA_norm"] = calls["geneA"].astype(str).apply(lambda value:
";".join(sorted({normalize_id(token) for token in value.split(";")})))
    calls["geneB_norm"] = calls["geneB"].astype(str).apply(lambda value:
";".join(sorted({normalize_id(token) for token in value.split(";")})))
    expanded = []
    for _, row in calls.iterrows():
        expanded.extend(expand_rows(row))
    calls_expanded = pd.DataFrame(expanded)
    truth = pd.read_csv(args.truth_tsv, sep="\t").rename(columns={"cdsA_id": "geneA", "cdsB_id":
"geneB"})
    truth["geneA"] = truth["geneA"].astype(str)
    truth["geneB"] = truth["geneB"].astype(str)
    if args.unordered:
        calls_expanded["pair"] = calls_expanded.apply(lambda rec: "|".join(sorted([rec["geneA"],
rec["geneB"]])), axis=1)
        truth["pair"] = truth.apply(lambda rec: "|".join(sorted([rec["geneA"], rec["geneB"]])),
axis=1)
    else:
        calls_expanded["pair"] = calls_expanded["geneA"] + "|" + calls_expanded["geneB"]
        truth["pair"] = truth["geneA"] + "|" + truth["geneB"]
    truth_pairs = set(truth["pair"])
    call_pairs = set(calls_expanded["pair"])
    tp_pairs = call_pairs & truth_pairs
    fp_pairs = call_pairs - truth_pairs
    fn_pairs = truth_pairs - call_pairs
    tp, fp, fn, precision, recall, f1 = compute_metrics(tp_pairs, fp_pairs, fn_pairs)
    ensure_parent_dir(args.out_prefix)
    with open(f"{args.out_prefix}.metrics.tsv", "w") as handle:
        handle.write("TP\tFP\tFN\tprecision\trecall\tF1\n")
        handle.write(f"{tp}\t{fp}\t{fn}\t{precision:.4f}\t{recall:.4f}\t{f1:.4f}\n")
    calls_expanded[calls_expanded["pair"].isin(tp_pairs)].to_csv(f"{args.out_prefix}.tp.tsv",
sep="\t", index=False)
    calls_expanded[calls_expanded["pair"].isin(fp_pairs)].to_csv(f"{args.out_prefix}.fp.tsv",
sep="\t", index=False)
    truth[truth["pair"].isin(fn_pairs)].to_csv(f"{args.out_prefix}.fn.tsv", sep="\t", index=False)
if __name__ == "__main__":
    main()
```
